# Animal models of chemotherapy-induced peripheral neuropathy: a machine-assisted systematic review and meta-analysis: A comprehensive summary of the field to inform robust experimental design

**DOI:** 10.1101/293480

**Authors:** Gillian L. Currie, Helena N. Angel-Scott, Lesley Colvin, Fala Cramond, Kaitlyn Hair, Laila Khandoker, Jing Liao, Malcolm Macleod, Sarah K. McCann, Rosie Morland, Nicki Sherratt, Robert Stewart, Ezgi Tanriver-Ayder, James Thomas, Qianying Wang, Rachel Wodarski, Ran Xiong, Andrew S.C. Rice, Emily Sena

**Affiliations:** Centre for Clinical Brain Sciences, University of Edinburgh London; Pain Research, Dept Surgery and Cancer, Imperial College London; Department of Anaesthesia, Critical Care & Pain; Centre for Clinical Brain Sciences, University of Edinburgh; FRCP, Centre for Clinical Brain Sciences, University of Edinburgh; Institute of Education, London

## Abstract

**Background and aims:** Chemotherapy-induced peripheral neuropathy (CIPN) can be a severely disabling side-effect of commonly used cancer chemotherapeutics, requiring cessation or dose reduction, impacting on survival and quality of life. Our aim was to conduct a systematic review and meta-analysis of research using animal models of CIPN to inform robust experimental design.

**Methods:** We systematically searched 5 online databases (PubMed, Web of Science, Citation Index, Biosis Previews and Embase (September 2012) to identify publications reporting *in vivo* CIPN modelling. Due to the number of publications and high accrual rate of new studies, we ran an updated search November 2015, using machine-learning and text mining to identify relevant studies.

All data were abstracted by two independent reviewers. For each comparison we calculated a standardised mean difference effect size then combined effects in a random effects meta-analysis. The impact of study design factors and reporting of measures to reduce the risk of bias was assessed. We ran power analysis for the most commonly reported behavioural tests.

**Results:** 341 publications were included. The majority (84%) of studies reported using male animals to model CIPN; the most commonly reported strain was Sprague Dawley rat. In modelling experiments, Vincristine was associated with the greatest increase in pain-related behaviour (−3.22 SD [−3.88; −2.56], n=152, p=0). The most commonly reported outcome measure was evoked limb withdrawal to mechanical monofilaments. Pain-related complex behaviours were rarely reported. The number of animals required to obtain 80% power with a significance level of 0.05 varied substantially across behavioural tests. Overall, studies were at moderate risk of bias, with modest reporting of measures to reduce the risk of bias.

**Conclusions:** Here we provide a comprehensive summary of the field of animal models of CIPN and inform robust experimental design by highlighting measures to increase the internal and external validity of studies using animal models of CIPN. Power calculations and other factors, such as clinical relevance, should inform the choice of outcome measure in study design.

## Introduction

Certain commonly used and effective cancer chemotherapeutic agents are neurotoxic to peripheral nerves. Treatment with these agents can result in distal symmetrical sensory polyneuropathy, and resultant dose limitation or reduction. Chemotherapy-induced peripheral neuropathy (CIPN) is a disabling side-effect, known to impair daily function and diminish quality of life [32]. Commonly used chemotherapeutic agents reported to cause neurotoxic effects include platinum derivatives, taxanes [40], vinca alkaloids, epothilones and newer agents (e.g thalidomide and bortezomib) [48]. The predominant sensory phenotype in patients treated with oxaliplatin or docetaxel is sensory loss affecting both upper and lower extremities. This includes bilateral paraesthesiae, often described as numbness or tingling. Other less common sensory symptoms include burning pain, pricking pain and cold allodynia [44]. CIPN can present clinically in two distinct forms: an acute, chemotherapy dose-related, and often dose-limiting, polyneuropathy, which in many cases resolves in many patients once the chemotherapy ceases. In some patients this will persist, with other patients developing symptoms after treatment has finished; and a chronic, often painful, distal sensory neuropathy is still present in 33% of patients one year after completion of treatment [39].

The pathogenesis of CIPN is thought to involve diverse mechanisms depending on the chemotherapeutic agent used. These mechanisms include chemotoxicity in the dorsal root ganglion (DRG) and DNA within the cell body of the DRG, alterations to transmembrane receptors and channels, interference with microtubular structure, damage to mitochondria, myelin degeneration and modulation of ion channels [2]. It is proposed that such mechanisms lead to the “dying-back” axonal degeneration characteristic of many sensory polyneuropathies.

Animal models of CIPN are used to investigate the pathophysiology of CIPN, and test potential therapies [19]. Administration of the chemotherapeutic agents can lead to the development of a sensory neuropathy, and behavioural tests are used to assess the sensory symptoms such as evoked pain and locomotor activity. Behavioural tests are less commonly used to attempt to measure sensory paraesthesia observed in patients [19].

Systematic review involves systematically locating and appraising all evidence relevant to a pre-defined research question, to provide a complete and unbiased summary of available evidence, and can be an invaluable tool to help make sense of the vast scientific literature.

Our aim was to provide a systematic overview of research in the field of *in vivo* animal modelling of CIPN, with a focus on the reporting pain-related behavioural outcome measures. The ultimate aim was to provide useful information for preclinical researchers to guide improvements in *in vivo* modelling.

## Methods

This review forms part of a larger review of all *in vivo* models of neuropathic pain and the full protocol can be found here: (www.dcn.ed.ac.uk/camarades/research.html#protocols). All methods were pre-specified in a study protocol.

### Search Strategy

We originally systematically searched 5 online databases (PubMed, Web of Science, Biosis Citation Index, Biosis Previews and Embase) in September 2012 to identify publications reporting *in vivo* modelling of CIPN, and the use of a pain-related behavioural outcome measure was reported. The search terms used for each database are detailed in Appendix 1. Search results were limited to animal studies using search filters, adapted for use in all databases [11; 22]. Because of the large number of publications and high accrual rate of publication of new studies, we ran an updated search in November 2015, using machine-learning and text-mining. This updated search included 4 online databases (PubMed, Web of Science, Bisosis Citation Index and Embase) and used an updated animal filter [10]. Biosis Previews was no longer available.

### Machine-learning and text mining

We used machine-learning to facilitate the screening of publications reporting animal models of CIPN.

The screening stage of a systematic review involves categorising records identified from the search as either ‘Included’ or ‘Excluded’ in the systematic review, and was performed by two independent reviewers. In our original search of 33,184 unique publications, the screening stage took 18 person months in total. The publications from our original search (with included/excluded decision based on initial screening) were used as a training set for machine-learning approaches: 13 classifiers were created and applied to the updated search (11,880 unique publications). We used the metrics of sensitivity (criteria set at: >95%) and specificity to evaluate the performance of machine-learning and used these measures to choose the classifier that performed best for our data set. Sensitivity was defined as the proportion of included publications that were correctly identified by the machine-learning algorithm, and specificity as the proportion of excluded publications that were correctly identified by the machine-learning algorithm (human reviewers’ final decision taken as correct). A 10% random sample of the machine-learning decisions was generated using a random number generator and the unique publication identifying numbers for the updated search, were checked for inclusion/exclusion by two independent investigators. Text mining was used on the publications identified by machine-learning to identify studies reporting animal models of CIPN. Briefly, this involved searching for specific chemotherapy terms within the title and abstract of the identified publications, which were then screened for inclusion by two independent reviewers.

### Inclusion and exclusion criteria

Studies that reported the use of animal models of neuropathy induced by chemotherapeutic agent administration and testing for a pain-related behavioural outcome measure, characterisation of a model or pharmacological intervention studies that report a pain-related behavioural outcome measure with a suitable control, and the number of animals per group and the mean and its variance (standard error of the mean (SEM) or standard deviation (SD)), were included in this analysis.

Studies that did not report testing for a pain-related behavioural outcome measure or that reported administration of an intervention before model induction, administration of co-treatments, transgenic studies or *in vitro* studies were excluded from this analysis.

### Measures to reduce the risk of bias and measures of reporting

We assessed methodological quality by looking at reporting of 5 measures to reduce the risk of bias; blinded assessment of outcome, random allocation to group, allocation concealment, animal exclusions and a sample size calculation. We also assessed the reporting of statement of potential conflicts of interest and of compliance with animal welfare regulations [27; 29].

### Data abstraction

Data were abstracted to the CAMARADES Data Manager (Microsoft Access). For all included studies, we abstracted details of; publication, animal husbandry, animal species and strain, model, intervention, and other experiment details (Table 1). Data presented graphically was abstracted using digital ruler software (Universal Desktop Ruler, AVPSoft.com or Adobe ruler) to determine values. Where multiple time points were presented we abstracted the time point that showed the greatest difference between model and control groups, or the greatest difference between treatment and control groups. If the type of variance (e.g. SEM or SD) was not reported it was recorded as SEM as this provides a more conservative estimate and avoids the study being given undue weight in the meta-analysis. All data were abstracted by two independent reviewers.

**Table 1.** Data extracted from each publication.

| Publication level |  |  |  |
| --- | --- | --- | --- |
| Publication | Husbandry |  |  |
| First author, corresponding author, year | Type of diet, type of bedding, enrichment, habituation time pre-study, cage cleaning frequency, number per cage, whether neuropathic pain models were housed in the same cages as Sham animals, number of hours in light cycle, room temperature, humidity and food availability |  |  |
| Experiment level |  |  |  |
| Animal | Model | Intervention | Experiment |
| Species, strain, sex, supplier, country of supplier | Chemotherapeutic agent, dose, number of doses, route of administration | Time of drug administration, dose, route of administration | Time of assessment, number of animals per group, mean outcome, variance, number of treatment groups per control |

### Data reconciliation

Publication level data abstracted by two independent reviewers were compared and any discrepancies were checked and corrected. Outcome level data were compared and discrepancies were checked and corrected. Individual comparison effect sizes were calculated for both reviewer’s data sets and those that differed by ≥10% were checked and corrected. Where individual comparisons differed by <10% we took a mean of the two effect sizes and errors (SEM/SD).

### Data analysis

We assessed the impact of animal species in stratified meta-analysis. We specified *a priori* that if species accounted for a significant proportion of heterogeneity each species would be analysed separately. If species did not account for a significant proportion of heterogeneity then all species (e.g. mouse and rat) would be analysed together.

We analysed all behavioural outcome measures reported. Behavioural outcome measures were split into pain-related and other behavioural outcome measures, and then grouped by subtype of outcome measure (full list of behavioural outcome measures and behavioural tests can be found in Tables 2 and 3).

**Table 2.**
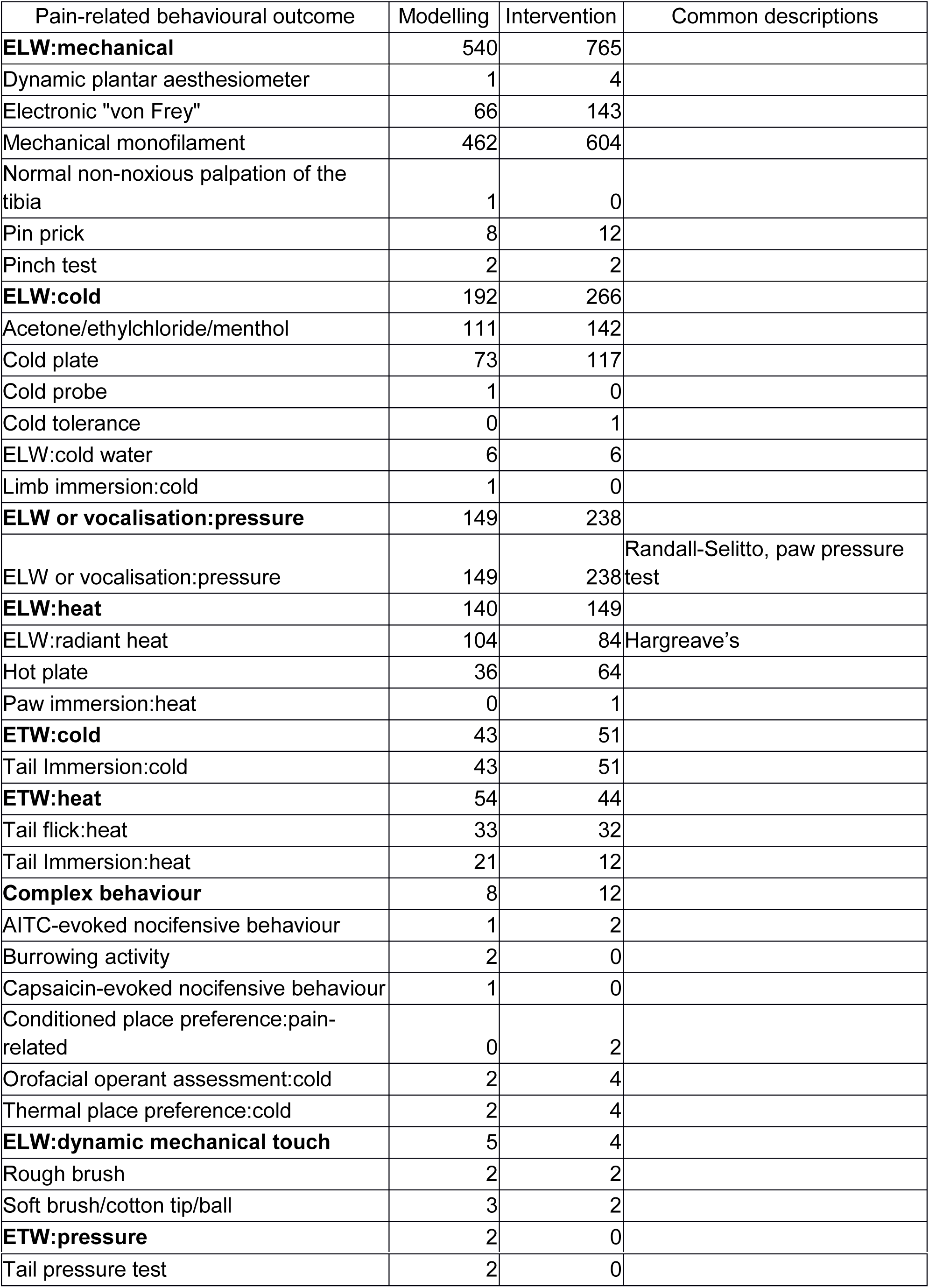
Pain-related behavioural outcome measures across intervention and modelling experiments. Numbers indicate the number of individual comparisons. AITC, TRPA1 agonist allyl isothiocyanate; ELW, Evoked limb withdrawal (ELW);, ETW, evoked tail withdrawal.

| Pain-related behavioural outcome | Modelling | Intervention | Common descriptions |
| --- | --- | --- | --- |
| **ELW:mechanical** | 540 | 765 |  |
| Dynamic plantar aesthesiometer | 1 | 4 |  |
| Electronic "von Frey" | 66 | 143 |  |
| Mechanical monofilament | 462 | 604 |  |
| Normal non-noxious palpation of the tibia | 1 | 0 |  |
| Pin prick | 8 | 12 |  |
| Pinch test | 2 | 2 |  |
| **ELW:cold** | 192 | 266 |  |
| Acetone/ethylchloride/menthol | 111 | 142 |  |
| Cold plate | 73 | 117 |  |
| Cold probe | 1 | 0 |  |
| Cold tolerance | 0 | 1 |  |
| ELW:cold water | 6 | 6 |  |
| Limb immersion:cold | 1 | 0 |  |
| **ELW or vocalisation:pressure** | 149 | 238 |  |
| ELW or vocalisation:pressure | 149 | 238 | Randall-Selitto, paw pressure test |
| **ELW:heat** | 140 | 149 |  |
| ELW:radiant heat | 104 | 84 | Hargreave's |
| Hot plate | 36 | 64 |  |
| Paw immersion:heat | 0 | 1 |  |
| **ETW:cold** | 43 | 51 |  |
| Tail Immersion:cold | 43 | 51 |  |
| **ETW:heat** | 54 | 44 |  |
| Tail flick:heat | 33 | 32 |  |
| Tail Immersion:heat | 21 | 12 |  |
| **Complex behaviour** | 8 | 12 |  |
| AITC-evoked nocifensive behaviour | 1 | 2 |  |
| Burrowing activity | 2 | 0 |  |
| Capsaicin-evoked nocifensive behaviour | 1 | 0 |  |
| Conditioned place preference:pain-related | 0 | 2 |  |
| Orofacial operant assessment:cold | 2 | 4 |  |
| Thermal place preference:cold | 2 | 4 |  |
| **ELW:dynamic mechanical touch** | 5 | 4 |  |
| Rough brush | 2 | 2 |  |
| Soft brush/cotton tip/ball | 3 | 2 |  |
| **ETW:pressure** | 2 | 0 |  |
| Tail pressure test | 2 | 0 |  |

**Table 3.** Other behavioural outcome measures across intervention and modelling experiments. Number of individual comparisons

| Other behavioural outcome | Modelling | Intervention |
| --- | --- | --- |
|  | Number of comparisons |  |
| **Locomotor function** | **73** | **37** |
| Rotarod:locomotor function | 34 | 23 |
| Grip test:motor strength | 14 | 1 |
| Locomotor activity | 10 | 6 |
| Open Field:distance travelled | 4 | 0 |
| Open field:exploratory behaviour | 2 | 2 |
| Open field:loading on hind limbs | 2 | 2 |
| Open field:immobility | 2 | 0 |
| Open Field:locomotor activity | 2 | 0 |
| Catwalk:gait alterations | 2 | 0 |
| Balance beam:motor coordination | 1 | 1 |
| Cannabinoid tetrad test | 0 | 1 |
| Cannabinoid tetrad test:motility | 0 | 1 |
| **Memory** | **8** | **0** |
| Novel location preference:preference score | 4 | 0 |
| Novel object preference:preference score | 4 | 0 |
| **Reward** | **4** | **6** |
| Conditioned place preference:rewarding effects | 4 | 6 |
| **Attention** | **3** | **0** |
| Pre-pulse inhibition:attention | 3 | 0 |

Pain-related outcome measures included evoked limb withdrawal to stimuli (mechanical stimuli/heat/cold/dynamic mechanical touch), evoked limb withdrawal or vocalisation to pressure stimuli, evoked tail withdrawal to stimuli (cold/heat/pressure), and complex behaviour). Other outcome measures included assessment of locomotor function, memory, reward, and attention. Behaviours were then nested by subtype for analysis. For each comparison we calculated a standardised mean difference (SMD) effect size, and combined these using a random effects meta-analysis with Restricted Maximum-Likelihood estimation of heterogeneity. When a single control group was used for multiple model or treatment groups, the impact was adjusted by dividing the number of animals in the control group by the number of treatment groups served. The Hartung and Knapp method is used to adjust test statistics and confidence intervals, which calculates the confidence intervals using: effect size + t(0.975,k-1)*standard error. The impact of study design factors (sex, species, therapeutic intervention, therapeutic intervention dose, time of administration, methods to induce the model including the chemotherapeutic agent, dose and route of administration and type of outcome measure) and study quality (as measured by reporting of measures to reduce bias and reporting measures) was assessed with stratified meta-analysis. The impact of time to assessment (defined as the interval between the first administration of chemotherapeutic agent and outcome measurement) and time to intervention administration (defined as the interval between the first administration of chemotherapeutic agent and the administration of intervention) were assessed using meta-regression. Stratified meta-analysis and meta-regression was performed using a Meta-analysis Online Platform (code available here: https://github.com/qianyingw/meta-analysis-app). A recent study from our group has indicated that using SMD estimates of effect sizes with stratified meta-analysis has a low statistical power to detect the effect of a variable of interest. This means that although we may not have sufficient power to detect an effect, we can have confidence that any significant results observed are true [47].

We applied a Bonferroni-Holmes correction as follows: study design in the modelling experiments (p<0.01), study design in the intervention experiments (p<0.007), the reporting of measures to reduce the risk of bias and of measures of reporting (p<0.007).

### Power analysis

To guide sample size estimation, we performed power calculations for the six most commonly reported behavioural tests. To do this, we ranked the observed mean difference effect sizes and pooled standard deviation, and used the 20th, 50th and 80th centile of each to calculate the number of animals required in hypothetical treatment and control groups. Calculations were based on the two sample two sided t-test, with 80% power and an alpha value of 0.05.

### Publication bias

Potential publication bias was assessed by visual inspection of funnel plots for asymmetry; Duval and Tweedie’s trim and fill analysis [12; 49] and Egger’s regression analysis [13] for small study effects were then performed. These analyses used modelling-individual comparisons and intervention-individual comparisons, as well as both pain-related and other outcome measures.

### Ranking of intervention efficacy

In a clinical systematic review of neuropathic pain [16], selected analgesic agents were ranked according efficacy, as measured by Numbers Needed to Treat (NNT) for 50 % pain relief. If preclinical studies included in this review reported use of these, or analogues of these agents, the interventions were ranked according to SMD effect size for attenuation of pain-related behaviour. The correlation between clinical and preclinical rank was assessed by Spearman’s rank correlation coefficient.

## Results

### Identification of Publications

Of the 33,184 unique publications using *in vivo* models of neuropathic pain that were identified by our original systematic search (September 2012), 181 were identified by independent reviewers as reporting CIPN, Figure 1.

**Figure 1.**
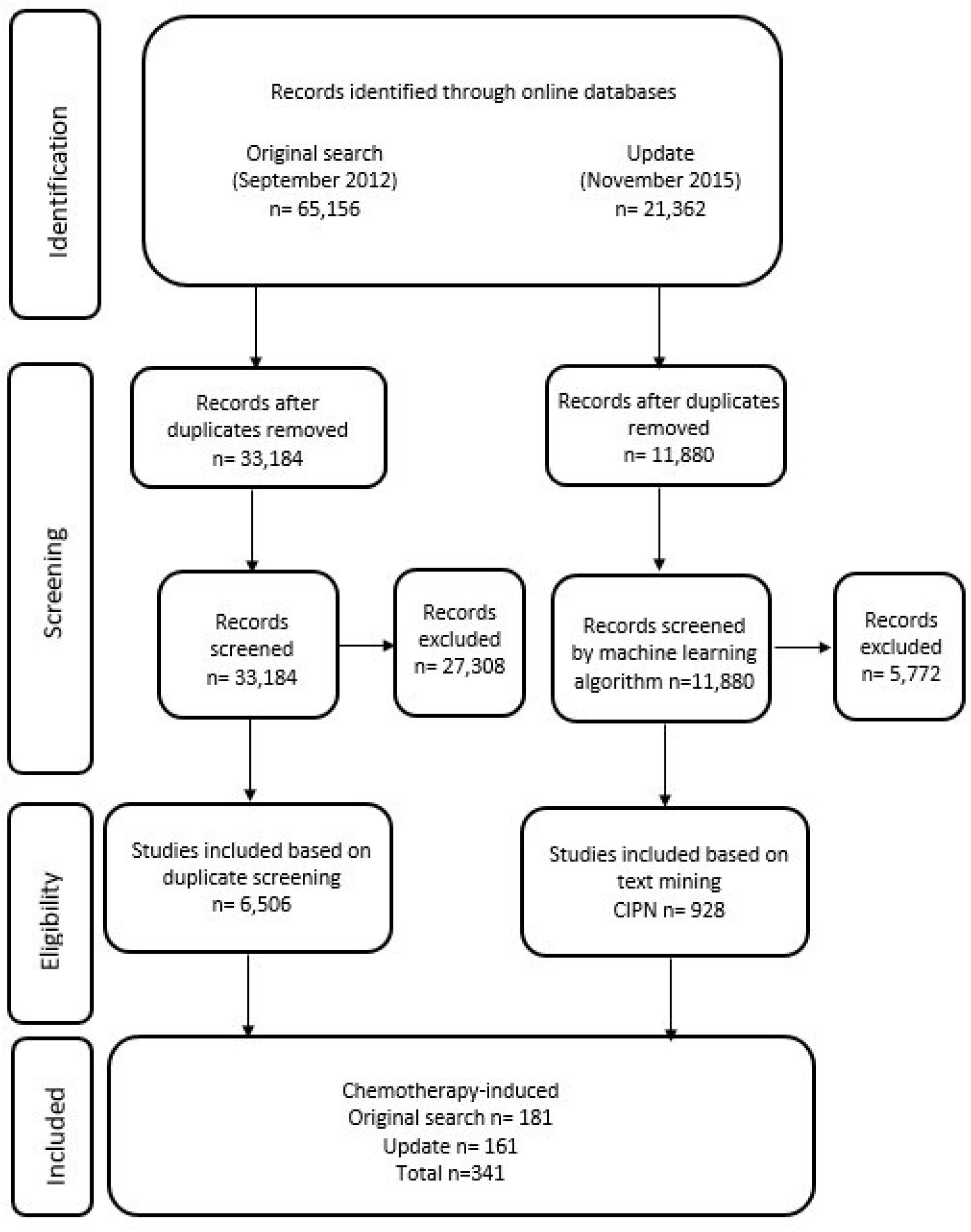
Flow diagram of included studies.

Of the 11,880 unique publications identified by the updated search (November 2015), 6,108 were identified using machine-learning as reporting CIPN. In the random 10% sample of screened publications (n = 1,188), the machine-learning approach with the best fit had a screening performance of 97% sensitivity 97%, 67% specificity, and 56% precision. Of the 359 studies identified by text mining as reporting animal models of CIPN, 160 met our inclusion criteria, as determined by two independent reviewers.

A total of 341 unique publications were identified from the two searches and included in this review. The rate of new publications per year are shown in Figure 2. Details of the 341 publications included in this study are available in Appendix 2.

**Figure 2.**
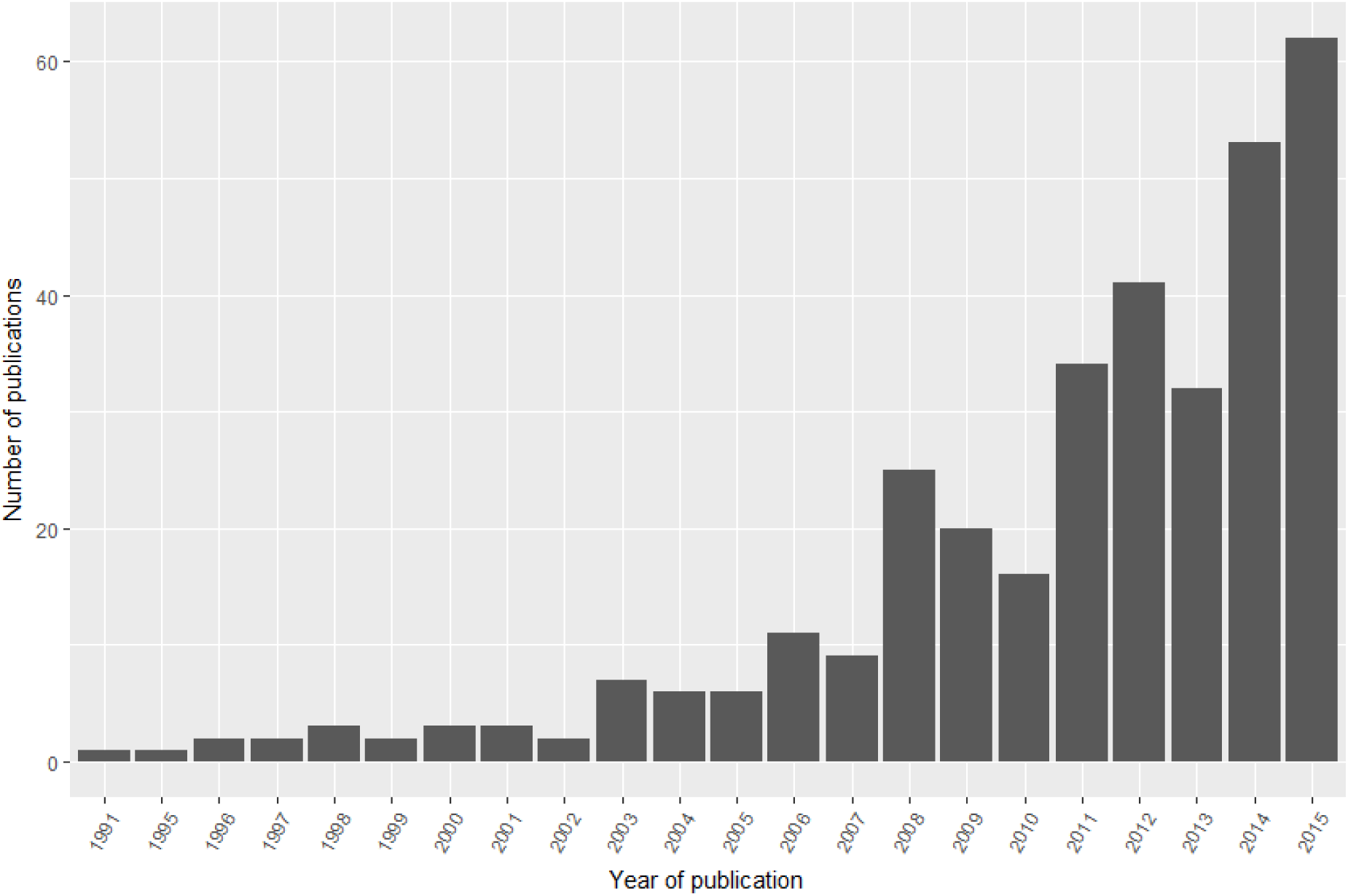
Number of included publications published each year.

### Outcome Measures

Across the 341 publications included, the most commonly reported pain-related outcome measure was evoked limb withdrawal to mechanical stimuli, most commonly assessed using mechanical monofilaments (Figure 3, table 2). The most frequently reported other behavioural outcome measures were those that assessed locomotor function, with rotarod apparatus used in the majorities of cases (Figure 4, table 3).

**Figure 3.**
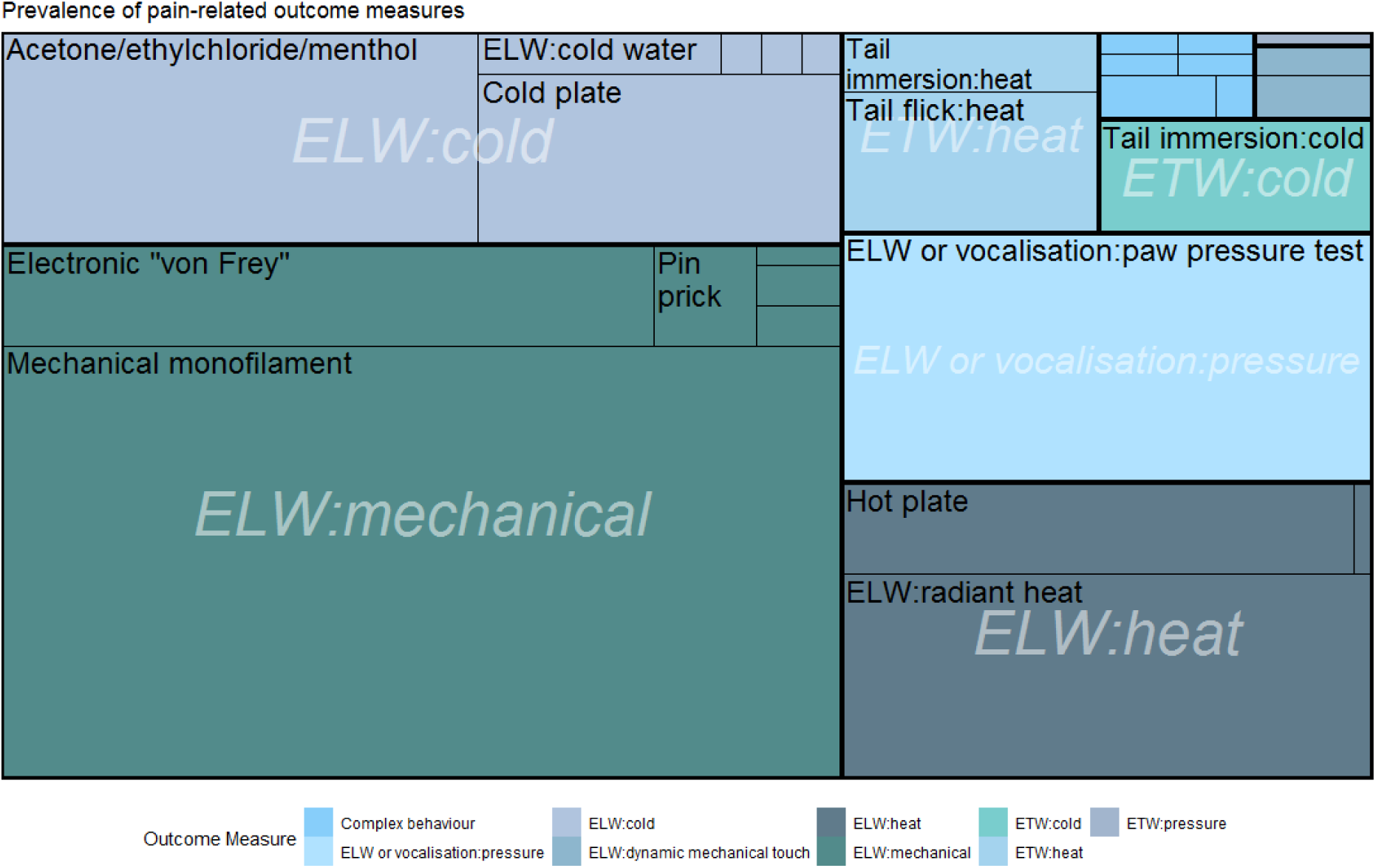
Tree plot of prevalence of pain-related outcome measures. Count of publications reporting each measure. ELW=evoked limb withdrawal, ETW=evoked tail withdrawal.

**Figure 4.**
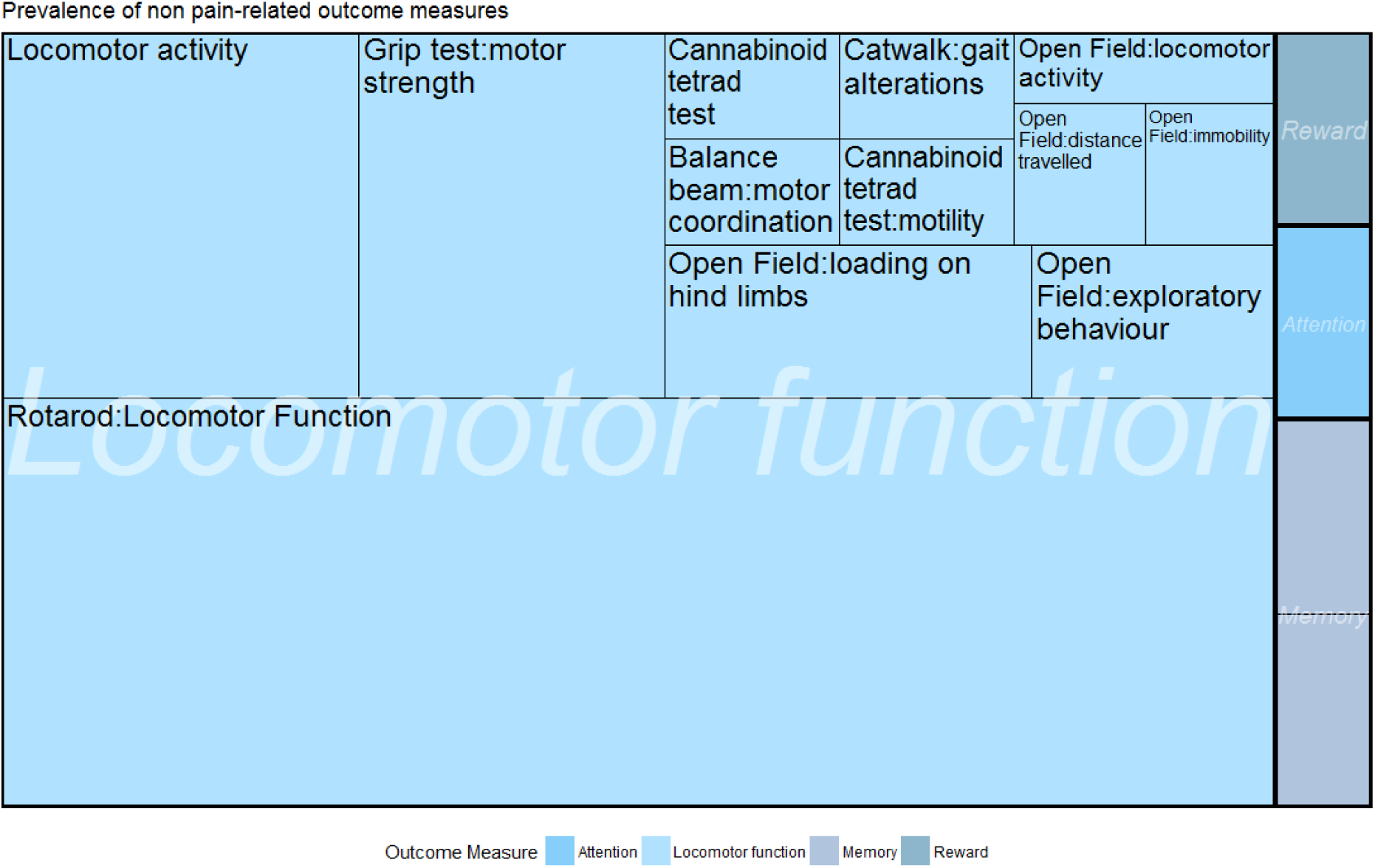
Tree plot of prevalence of other behavioural outcome measures, colours used to denote the same behavioural subtype. Count of publications reporting each measure.

### Interventions

A total of 307 different interventions were tested (Figure 5). The majority of interventions (80%) were only tested by one publication, and the most commonly reported interventions were gabapentin, morphine and pregabalin, reported in 26, 23 and 12 publications, respectively.

**Figure 5.**
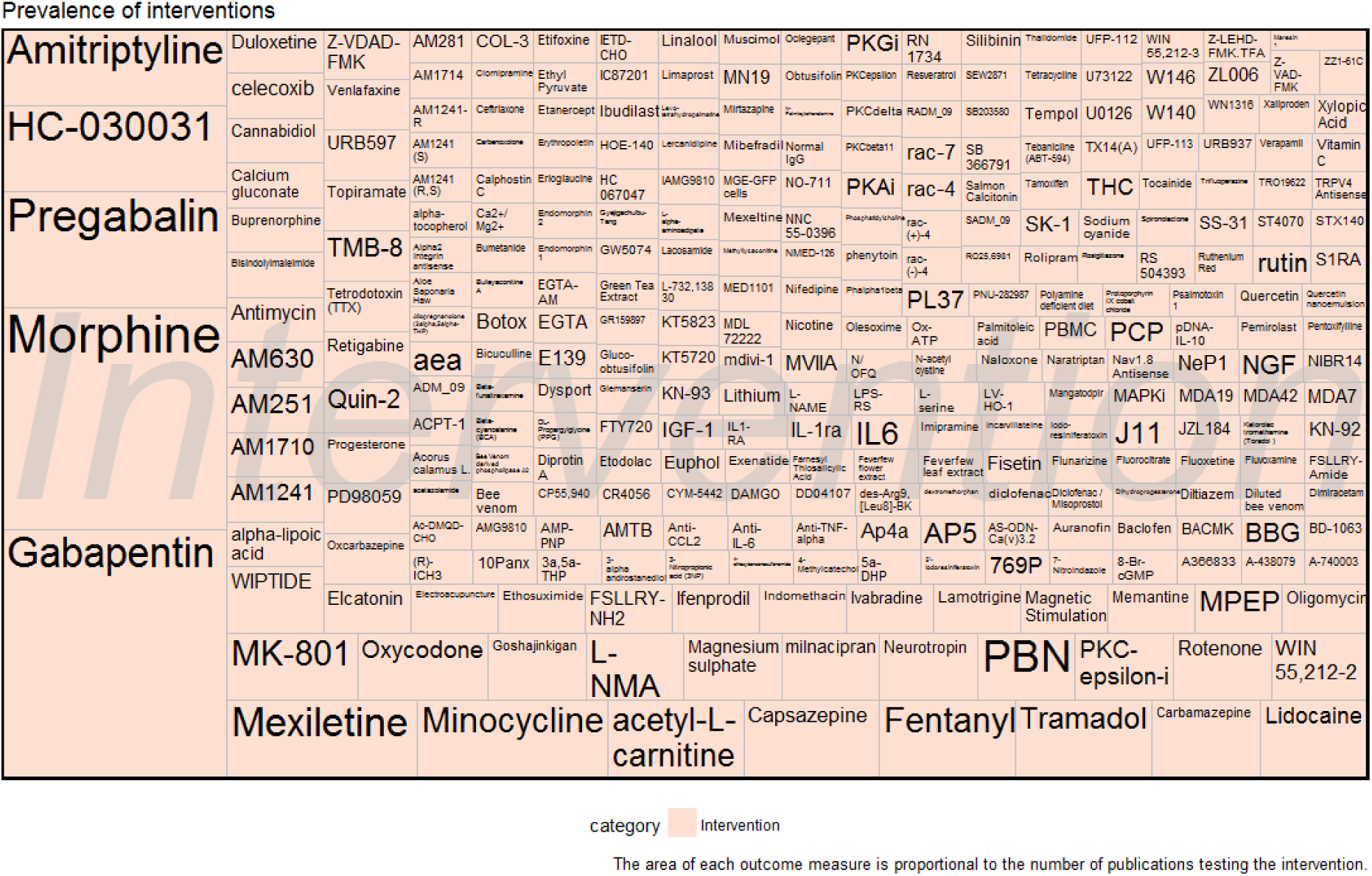
Tree plot of prevalence of interventions. 307 different interventions reported.

### Risk of bias

The reporting of measures to reduce the risk of bias was moderate across all included CIPN studies (n = 341): 51.3% (175) reported blinded assessment of outcome, 28.2% (96) reported randomisation to group, 17.6% (60) reported animal exclusions, 2.1% (7) reported the use of a sample size calculation, and 1.5% (5) reported allocation concealment. In terms of measures of reporting, 49.6% (168) of studies reported a conflict of interest statement and 96.8% (330), reported compliance with animal welfare regulation.

The details of methods used to implement randomisation and blinding and the methods and assumptions for sample size calculation were rarely reported: six publications reported that animals were randomly allocated to experimental groups using randomly generated number sequences and two publications reported this was done by block randomisation. One publication reported randomisation was performed by “picking animals randomly from a cage”, which is known not to be a valid form of randomisation [43]. Nine publications reported that blinded outcome assessment of outcome was achieved by using a different experimenter to perform assessment, and two publications reported that a group code was used. One study reported allocation concealment was achieved using a coded system.

Methods of sample size calculation were reported by five publications: three used published or previous results from the group and two performed a pilot study to inform sample size calculations.

## Modelling Experiments

### Animal Studies Modelling CIPN: Pain-Related Behavioural Outcome Measures

In modelling experiments using pain-related behavioural outcome measures, administration of chemotherapeutic agents led to an increase in pain-related behaviour compared to sham controls (−2.56 SD [-2.71; -2.41 95% CI], n=890 comparisons). The number of animals contributing to comparisons is shown in the figures (N). Species did not account for a significant proportion of the heterogeneity, and therefore mice and rat experiments were analysed together (mice: -2.63 SD [-2.87; -2.40 95% CI], n=344 comparisons; rats: -2.52 SD [-2.71; -2.32 95% CI], n=546 comparisons; p=0.27.

The most frequently reported outcome measure was evoked limb withdrawal to mechanical stimuli (540 individual comparisons), most commonly assessed using mechanical monofilaments to induce limb withdrawal (462 individual comparisons).

### Impact of study design

Pain-related outcome measure accounted for a significant proportion of the heterogeneity, with the evoked tail withdrawal to a pressure stimulus associated with the greatest increase in pain-related behaviour (−4.58 SD [-7.89;-1.27], n=2, p=0, Figure 6a). The most commonly reported pain-related outcome was evoked limb withdrawal to a mechanical stimulus (n=404 comparisons, 6488 animals).

**Figure 6.**
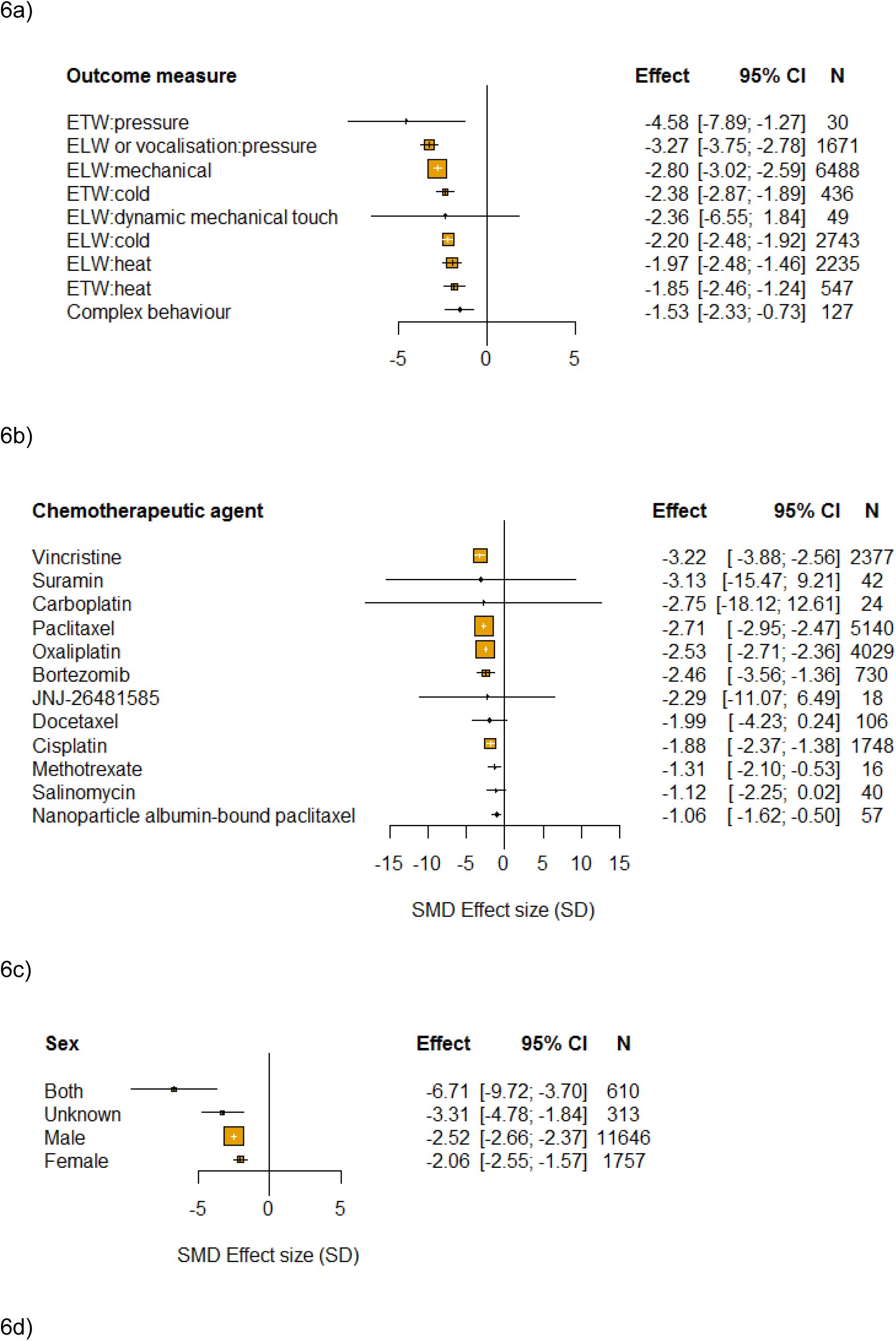
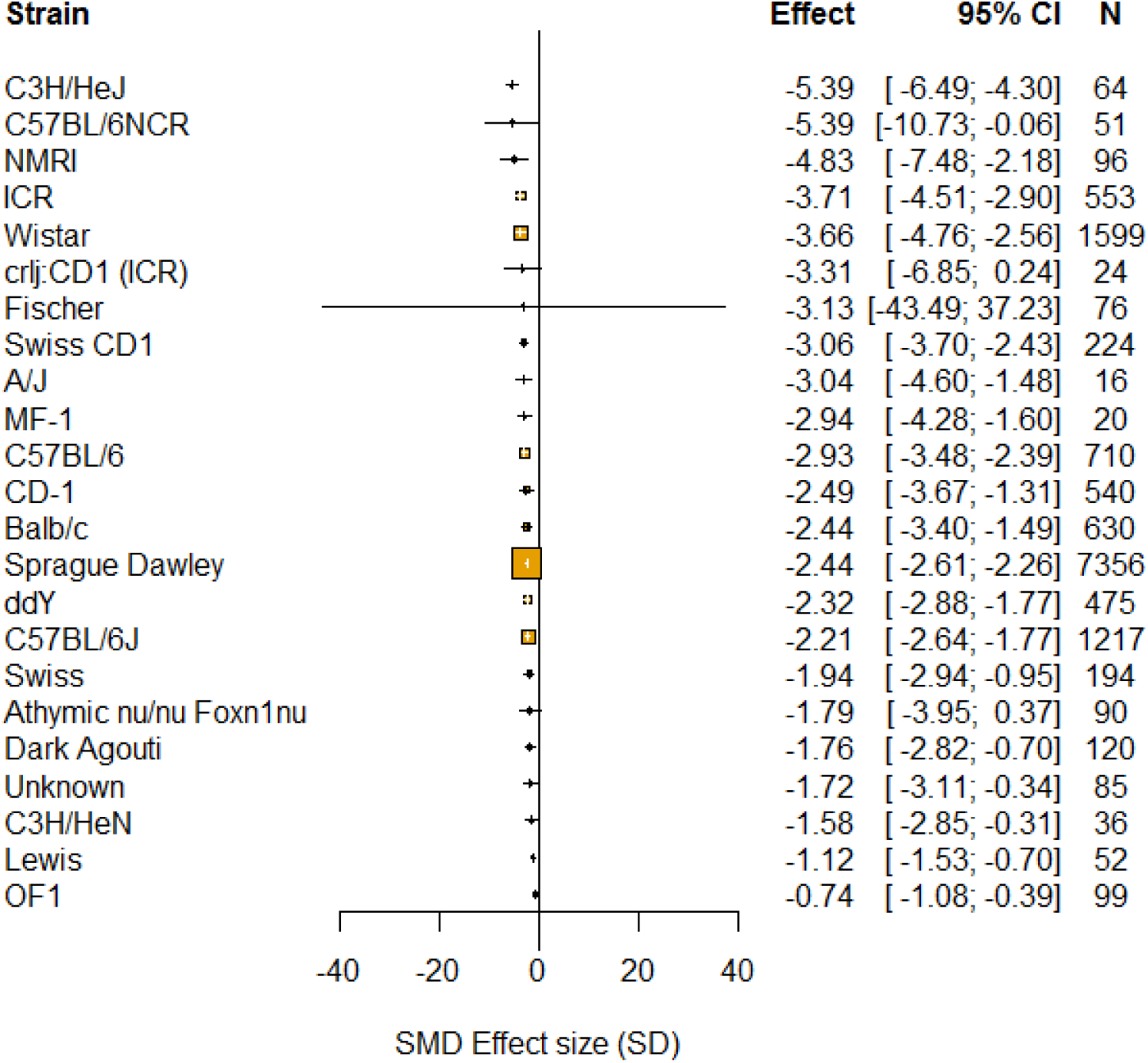
Impact of study design in modelling experiments reporting pain-related behavioural outcomes. The size of the squares represents the number of nested comparisons that contribute to that data point and the value N represents the number of animals that contribute to that data point. a) Outcome measure accounted for a significant proportion of the heterogeneity. Evoked tail withdrawal (ETW), evoked limb withdrawal (ELW) and complex behaviours used to measure pain. b) Chemotherapeutic agent accounted for a significant proportion of the heterogeneity. c) Sex accounted for a significant proportion of the heterogeneity. d) Strain accounted for a significant proportion of the heterogeneity.

This systematic review identified twelve different chemotherapeutic agents used to model CIPN in animals (Table 4). The chemotherapeutic agent used accounted for a significant proportion of the heterogeneity observed (p=0). Vincristine was associated with the greatest increase in pain-related behaviour (Figure 6b).

**Table 4.** Model details including chemotherapeutic agents, route of administration and median cumulative dose and upper and lower quartiles.

| Animal | Type | Chemotherapeutic agent | Route of administration | Median | Q1 | Q3 |
| --- | --- | --- | --- | --- | --- | --- |
| Mouse | Hydroxamate histone | JNJ-26481585 | Subcutaneous | 32.5 | 23.75 | 41.25 |
|  | Other | Salinomycin | Intraperitoneal | 140 | 140 | 140 |
|  | Platinum compounds | Cisplatin | Hindpaw | 0.004 | 0.0022 | 0.022 |
|  |  |  | Intraperitoneal | 20.5 | 8.25 | 23 |
|  |  |  | Subcutaneous | 0.15 | 0.15 | 0.15 |
|  |  | Oxaliplatin | Hindpaw | 0.04 | 0.04 | 0.04 |
|  |  |  | Intraperitoneal | 10 | 3 | 30 |
|  |  |  | Intravenous | 23 | 13.5 | 31 |
|  |  |  | Subcutaneous | 0.04 | 0.04 | 0.04 |
|  | Proteasome inhibitors | Bortezomib | Intraperitoneal | 0.75 | 0.425 | 1.95 |
|  |  |  | Intravenous | 6.4 | 6.4 | 6.4 |
|  |  |  | Subcutaneous | 12 | 12 | 18 |
|  | Taxanes | Paclitaxel | Intraperitoneal | 10 | 5 | 16 |
|  |  |  | Intravenous | 190 | 165 | 230 |
|  |  |  | Subcutaneous | 10 | 10 | 10 |
|  |  |  | Tail vein | 75 | 75 | 75 |
|  | Vinca alkaloids | Vincristine | Intraperitoneal | 0.7 | 0.2 | 3.9 |
| Rat | Binding of growth factor | Suramin | Intravenous | 25 | 17.5 | 37.5 |
|  | Other | Methotrexate | Subcutaneous | 3.75 | 3.75 | 3.75 |
|  | Platinum compounds | Carboplatin | Intraperitoneal | 112.5 | 101.25 | 123.75 |
|  |  | Cisplatin | Intraperitoneal | 10 | 5 | 15 |
|  |  |  | Intravenous | 2 | 2 | 2 |
|  |  |  | Subcutaneous | 8 | 8 | 8 |
|  |  | Oxaliplatin | Intraperitoneal | 16 | 8 | 32 |
|  |  |  | Intravenous | 2 | 2 | 8 |
|  | Proteasome inhibitors | Bortezomib | Intraperitoneal | 0.7 | 0.6 | 1 |
|  |  |  | Intravenous | 4.8 | 4.2 | 4.8 |
|  | Taxanes | Docetaxel | Intravenous | 25 | 17.5 | 32.5 |
|  |  |  | Tail vein | 10 | 10 | 10 |
|  |  | Paclitaxel | Intraperitoneal | 8 | 7.25 | 8.5 |
|  |  |  | Intravenous | 24.5 | 9.75 | 33.25 |
|  |  |  | Tail vein | 8 | 8 | 8 |
|  |  | Nanoparticle | Intravenous | 26.8 | 25.4 | 28.2 |
|  | Vinca alkaloids | Vincristine | Intraperitoneal | 0.95 | 0.5 | 1 |
|  |  |  | Intravenous | 0.435 | 0.2125 | 0.75 |
|  |  |  | Subcutaneous | 0.42 | 0.42 | 0.42 |
|  |  |  | Tail | 0.625 | 0.5625 | 0.6875 |
|  | Taxanes<sup>a</sup> | Paclitaxel | Intrathecal | 6 | 3.1 | 13 |

<sup>a</sup>Drug doses are shown in mg/kg, except intrathecal Paclitaxel, shown in ng.

Sex accounted for a significant proportion of the heterogeneity (p=0). Studies that reported using both males and females reported an increase in pain-related behaviour compared to studies that did not report the sex of animals used or used only male or female animals (Figure 6c).

The time to assessment did not account for a significant proportion of heterogeneity (p=1).

A *post hoc* analysis found strain of animal accounted for a significant proportion of the heterogeneity(p=0, Figure 6d). The most commonly reported were Sprague Dawley rats - 2.44 SD [-2.61:-2.26 95% CI], n=439 comparisons.

### Animal Studies Modelling CIPN: Other Behavioural Outcomes

In modelling experiments using other behavioural outcomes (locomotor function, memory, reward behaviours, and attention), administration of chemotherapeutic agents led to a worsening in these behaviours compared to sham controls (−0.75 [-1.04; -0.47 95% CI], n=63 comparisons). Species did not account for a significant proportion of the heterogeneity (p=0.070), rats and mice were analysed together.

The most common other behavioural outcome reported in modelling experiments was locomotor function (73 comparisons); the most frequently reported assay for locomotor function was the rotarod (34 comparisons).

### Impact of study design

Outcome measure accounted for a significant proportion of the heterogeneity (p=2.11×10^−02^), with memory associated with the greatest worsening of condition (−0.96 SD [-1.65;-0.27 95% CI], Figure 7a).

**Figure 7.**
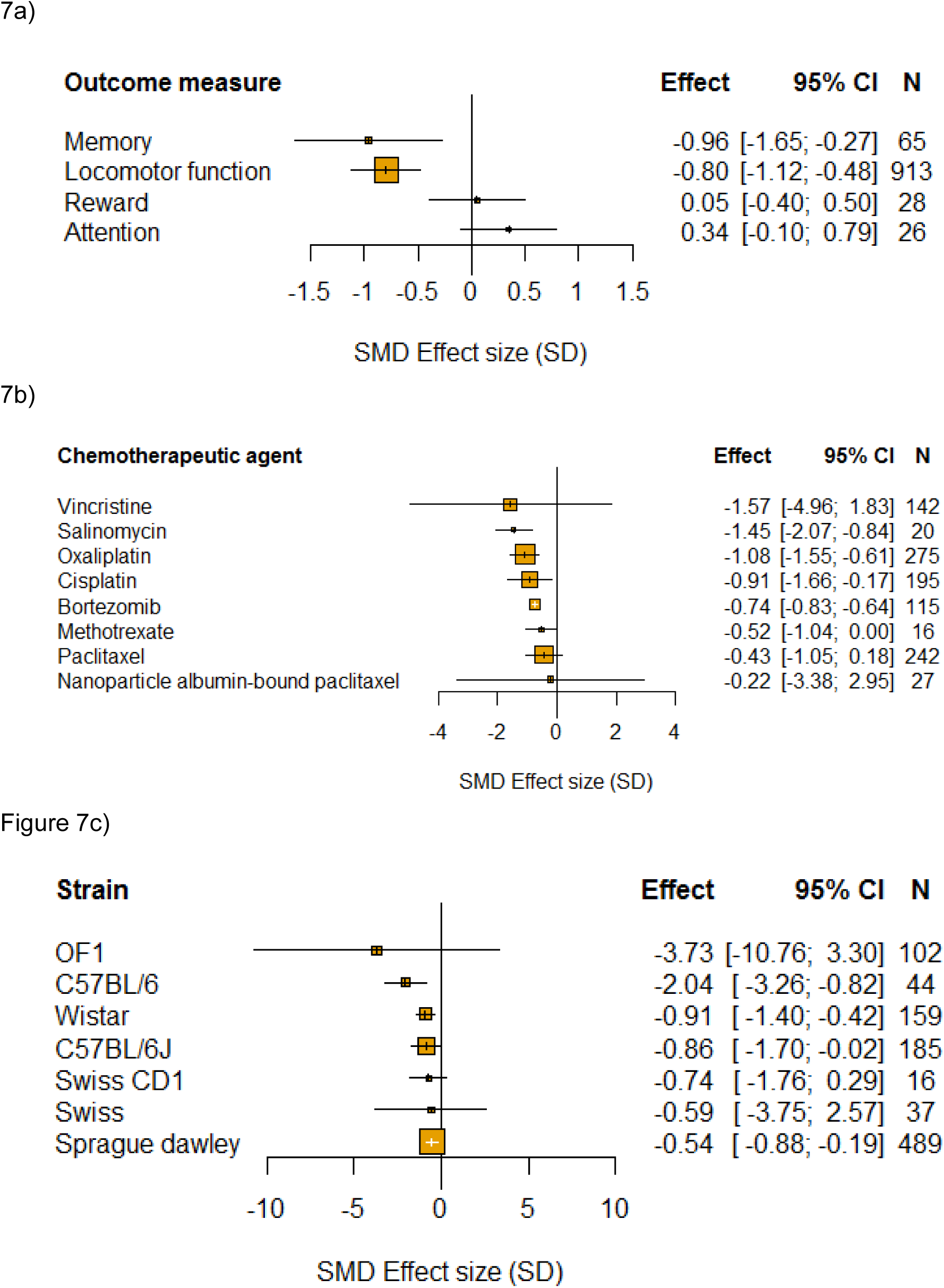
Impact of study design in modelling experiments using other behavioural outcomes. The size of the squares represents the number of nested comparisons that contribute to that data point and the value N represents the number of animals that contribute to that data point. a) Outcome measure accounted for a significant proportion of the heterogeneity. b) Chemotherapeutic agent accounted for a significant proportion of the heterogeneity. c) Strain accounted for a significant proportion of the heterogeneity.

Chemotherapeutic agent accounted for a significant proportion of the heterogeneity, (p=1.5×10^−04^), with vincristine associated with the greatest worsening in other behavioural outcomes (−1.57 SD [-4.96;1.83 95% CI], n=142 comparisons, Figure 7b).

Sex (p=0.70) and time to intervention administration (p=1.00) did not account for a significant proportion of the heterogeneity in modelling experiments using other behavioural outcomes.

A *post hoc* analysis found that strain accounted for a significant proportion of the heterogeneity (p=5.3×10^−04^, Figure 7c).

### Power analysis

The number of animals required to obtain an 80% power with a significance level of 0.05 varied substantially across the behavioural tests. For each of mechanical monofilaments, Randall-Selitto paw pressure test, Electronic "von Frey", acetone test/ethylchloride spray, cold plate, and Plantar Test (Hargreave’s method), we calculated the number of animals required in both model and sham groups.

When both mean difference effect sizes and pooled standard deviation were the 50% centile, the number of animals required ranged from 3 (Randall-Selitto and electronic “von Frey”/cold plate) to 7 (Hargreave’s) (Figure 8). Keeping the mean difference effect size in the 50% centile and increasing pooled standard deviation to the 80% centile increased the number of animals required per group across all behavioural tests, from 10 (electronic “von Frey” and cold plate) to 30 (acetone test/ethylchloride spray) animals. Reducing the mean difference effect size to the 20% centile and pooled standard deviation to the 50% centile, also increases the number of animals required, from 4 (Randall-Selitto) to 98 (mechanical monofilaments) (Figure 8). The values for the 20th, 50th and 80th centile of Mean Difference effect sizes and standard deviations for each behavioural test are available in supplementary Appendix 1.

**Figure 8.**
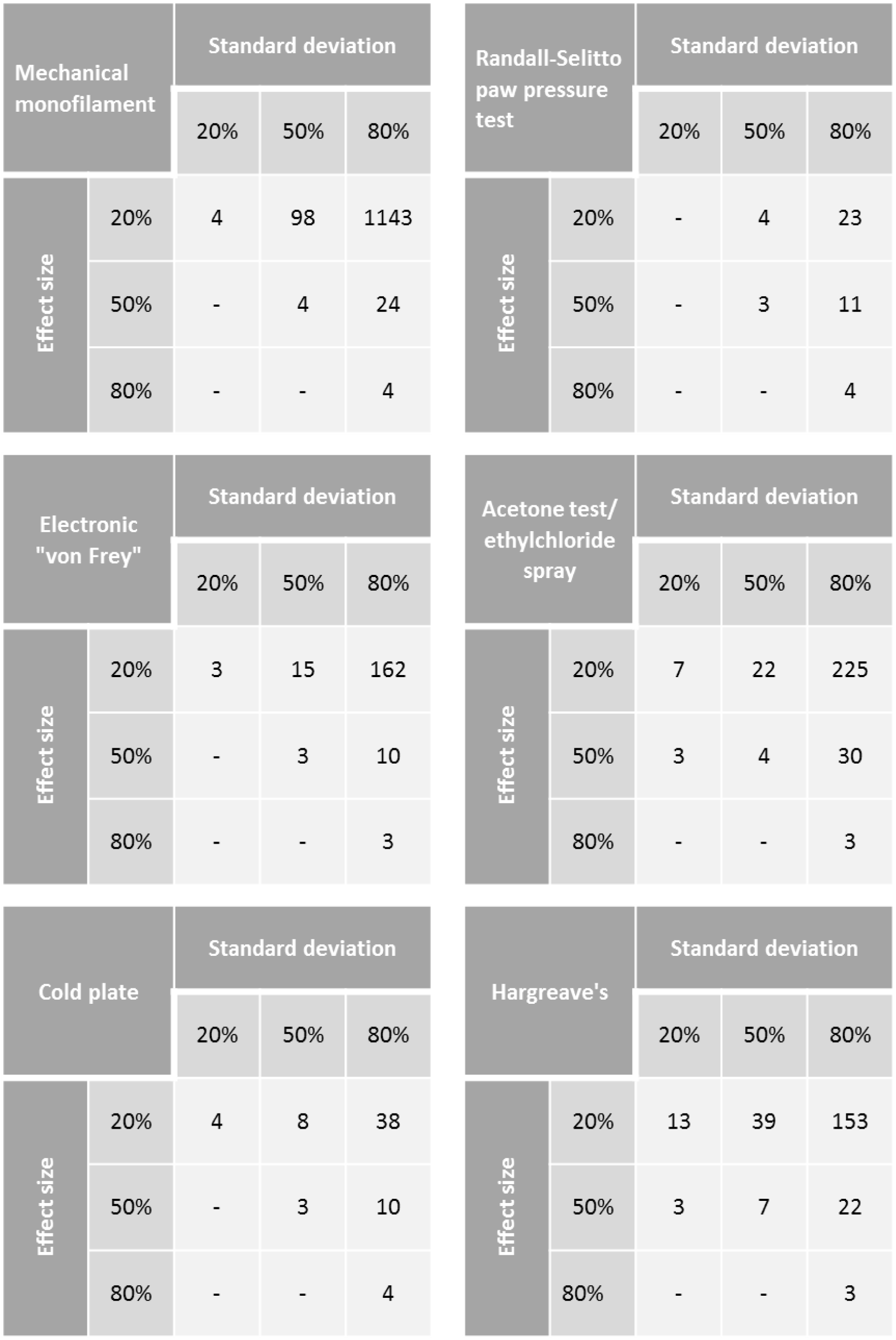
Power analysis for modelling experiments. Number of animals required per group to obtain 80% power with a significance level of 0.05 using mechanical monofilaments, Randall-Selitto paw pressure test, Electronic “von Frey”, Acetone test/ethylchloride spray, cold plate and Hargreave’s. Effect sizes calculated by Mean Difference. Blank squares could not be calculated.

### Animal Studies Modelling CIPN: Impact of Measures to Reduce the Risk of Bias and Measures of Reporting

In CIPN modelling experiments involving a pain-related behavioural outcome measure, reporting of blinded assessment of outcome (p=0) and animal exclusions accounted for a significant proportion of the heterogeneity (p=6×10^−08^), with failure to report these measures associated with greater estimates of effect. Reporting of randomisation, allocation concealment and sample size calculation did not account for a significant proportion of the heterogeneity (Figure 9).

**Figure 9.**
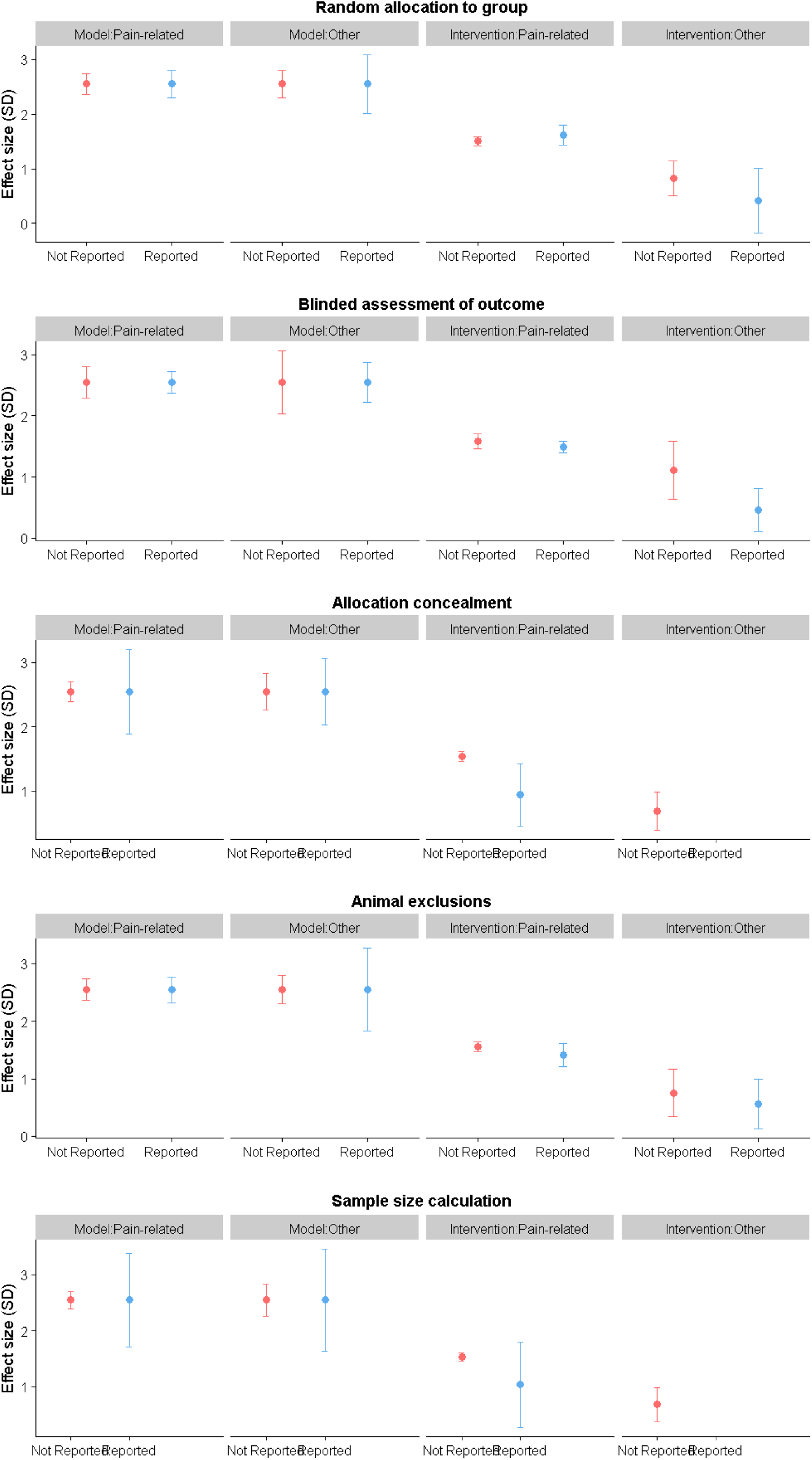
Effect sizes associated with measures to reduce the risk of bias in modelling and intervention experiments. Reporting of; a) random allocation to group b) blinded assessment of outcome c) allocation concealment d) animal exclusions and e) sample size calculations.

Regarding reporting measures, compliance with animal welfare regulations accounted for a significant proportion of the heterogeneity (p=9.9×10^−06^), with failure to report this was associated with a decreased estimate of effect (Figure 10). However, reporting of a COI statement did not account for a significant proportion of the heterogeneity (Table 5, Figure 10).

**Table 5.** Modelling comparisons using pain-related behavioural outcome measures. Stratified meta-analysis results for reporting of measures to reduce the risk of bias measures and measures of reporting (p<0.007).

|  | Modelling comparisons pain-related (n=890) |  |  |  |  |  |
| --- | --- | --- | --- | --- | --- | --- |
|  | Accounts for a significant proportion of heterogeneity | Not reporting associated with | Reported | Effect | SE | No. of comparisons |
| Risk of bias |  |  |  |  |  |  |
| Blinded assessment of outcome | Yes | Increased estimate of effect | TRUE | -2.547 | 0.091 | 476 |
|  |  |  | FALSE | -2.601 | 0.132 | 414 |
| Random allocation to group | No | NA | TRUE | -2.561 | 0.126 | 272 |
|  |  |  | FALSE | -2.563 | 0.096 | 618 |
| Allocation concealment | No | NA | TRUE | -1.916 | 0.334 | 12 |
|  |  |  | FALSE | -2.571 | 0.078 | 878 |
| Sample size calculation | No | NA | TRUE | -2.384 | 0.428 | 15 |
|  |  |  | FALSE | -2.565 | 0.078 | 875 |
| Animal exclusions | Yes | Increased estimate of effect | TRUE | -1.946 | 0.113 | 189 |
|  |  |  | FALSE | -2.762 | 0.094 | 701 |
| Reporting |  |  |  |  |  |  |
| Compliance with animal welfare regulations | Yes | Decreased estimate of effect | TRUE | -2.58 | 0.078 | 872 |
|  |  |  | FALSE | -1.652 | 0.39 | 18 |
| Conflict of interest statement | No | NA | TRUE | -2.702 | 0.114 | 446 |
|  |  |  | FALSE | -2.421 | 0.102 | 444 |

**Figure 10.**
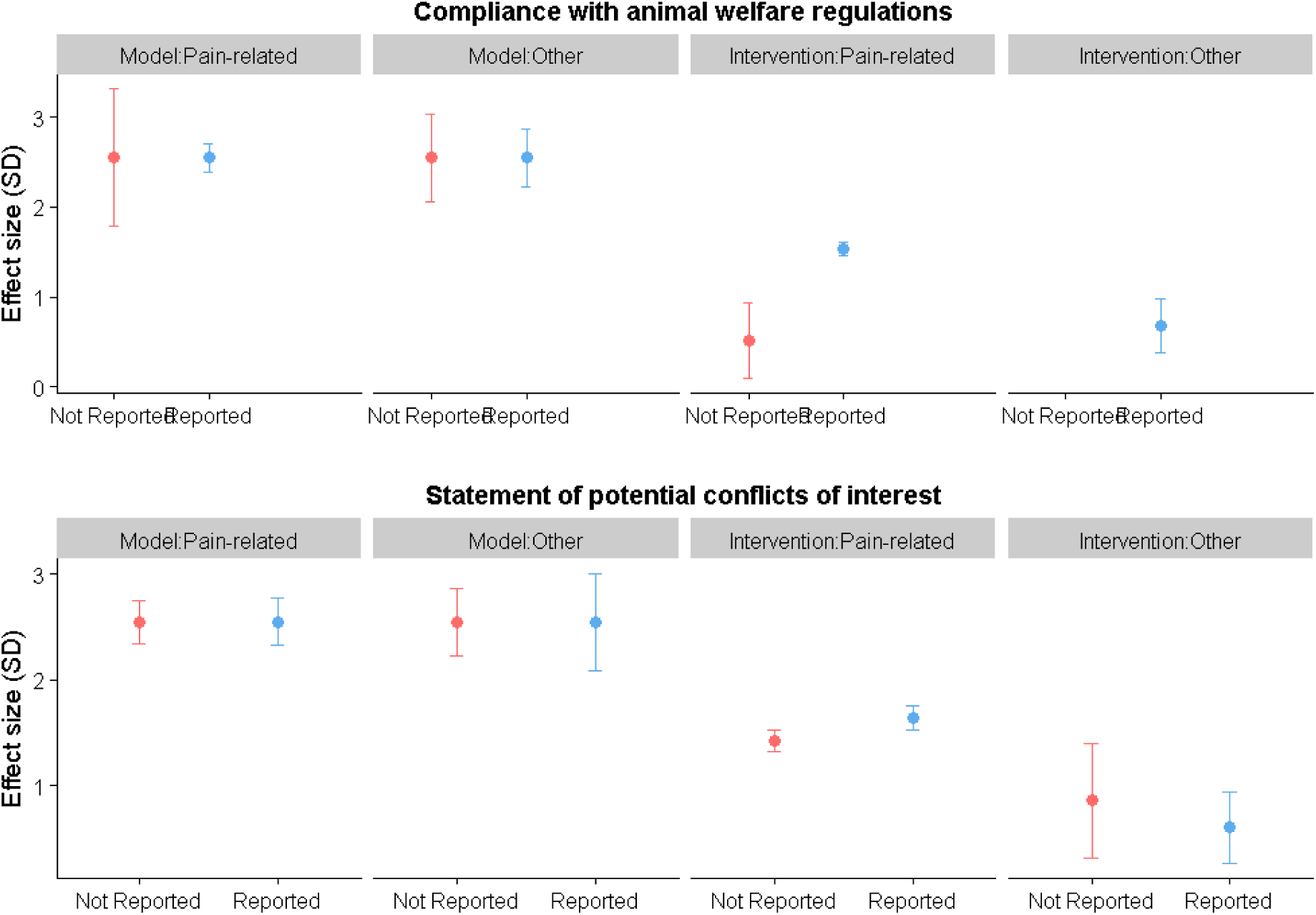
Effect sizes associated with measures of reporting in modelling and intervention experiments. Reporting of; a) compliance with animal welfare regulations and b) statement of potential conflict of interests.

In modelling experiments reporting other behavioural outcome measures, reporting of randomisation, blinding, allocation concealment, sample size calculation, or animal exclusions did not account for a significant proportion of the heterogeneity (Figure 9 and Table 6), nor did reporting of compliance with animal welfare regulations or a conflict of interest statement (Figure 10).

**Table 6.** Modelling comparisons using other behavioural outcome measures. Stratified meta-analysis results for reporting of measures to reduce the risk of bias and measures of reporting (p<0.007).

|  | Modelling comparisons other (n=63) |  |  |  |  |  |
| --- | --- | --- | --- | --- | --- | --- |
|  | Accounts for a significant proportion of heterogeneity | Not reporting associated with | Reported | Effect | SE | No. of comparisons |
| Risk of bias |  |  |  |  |  |  |
| Blinded assessment of outcome | No | NA | TRUE | -0.74 | 0.166 | 42 |
|  |  |  | FALSE | -0.76 | 0.261 | 21 |
| Random allocation to group | No | NA | TRUE | -1.036 | 0.271 | 24 |
|  |  |  | FALSE | -0.562 | 0.13 | 39 |
| Allocation concealment | No | NA | TRUE | -0.52 | 0.263 | 1 |
|  |  |  | FALSE | -0.762 | 0.146 | 62 |
| Sample size calculation | No | NA | TRUE | -0.155 | 0.462 | 6 |
|  |  |  | FALSE | -0.815 | 0.146 | 57 |
| Animal exclusions | No | NA | TRUE | -0.936 | 0.366 | 24 |
|  |  |  | FALSE | -0.641 | 0.124 | 39 |
| Reporting |  |  |  |  |  |  |
| Compliance with animal welfare regulations | No | NA | TRUE | -0.789 | 0.162 | 56 |
|  |  |  | FALSE | -0.556 | 0.251 | 7 |
| Conflict of interest statement | No | NA | TRUE | -0.764 | 0.232 | 28 |
|  |  |  | FALSE | -0.72 | 0.164 | 35 |

### Assessment of publication bias in animal studies modelling CIPN

For modelling experiments using pain-related behavioural outcome measures, there were 1133 individual comparisons (−2.59 SD [-2.72;-2.45 95%CI]). Visual inspection of funnel plots indicated asymmetry, suggesting theoretical missing studies (Figure 11a). Trim and fill analysis, imputed 318 theoretical missing studies on the right hand side of the funnel plot, resulting in a total of 1451 individual comparisons, (Figure 11b). Inclusion of these theoretical missing studies decreased the estimate of modelling-induced pain-related behaviour by 29% to -1.83 SD [-1.97;-1.68 95%CI]. Further, Egger’s regression line and 95% CI did not pass through the origin indicating small study effects, suggesting funnel plot asymmetry, Figure 11c.

**Figure 11.**
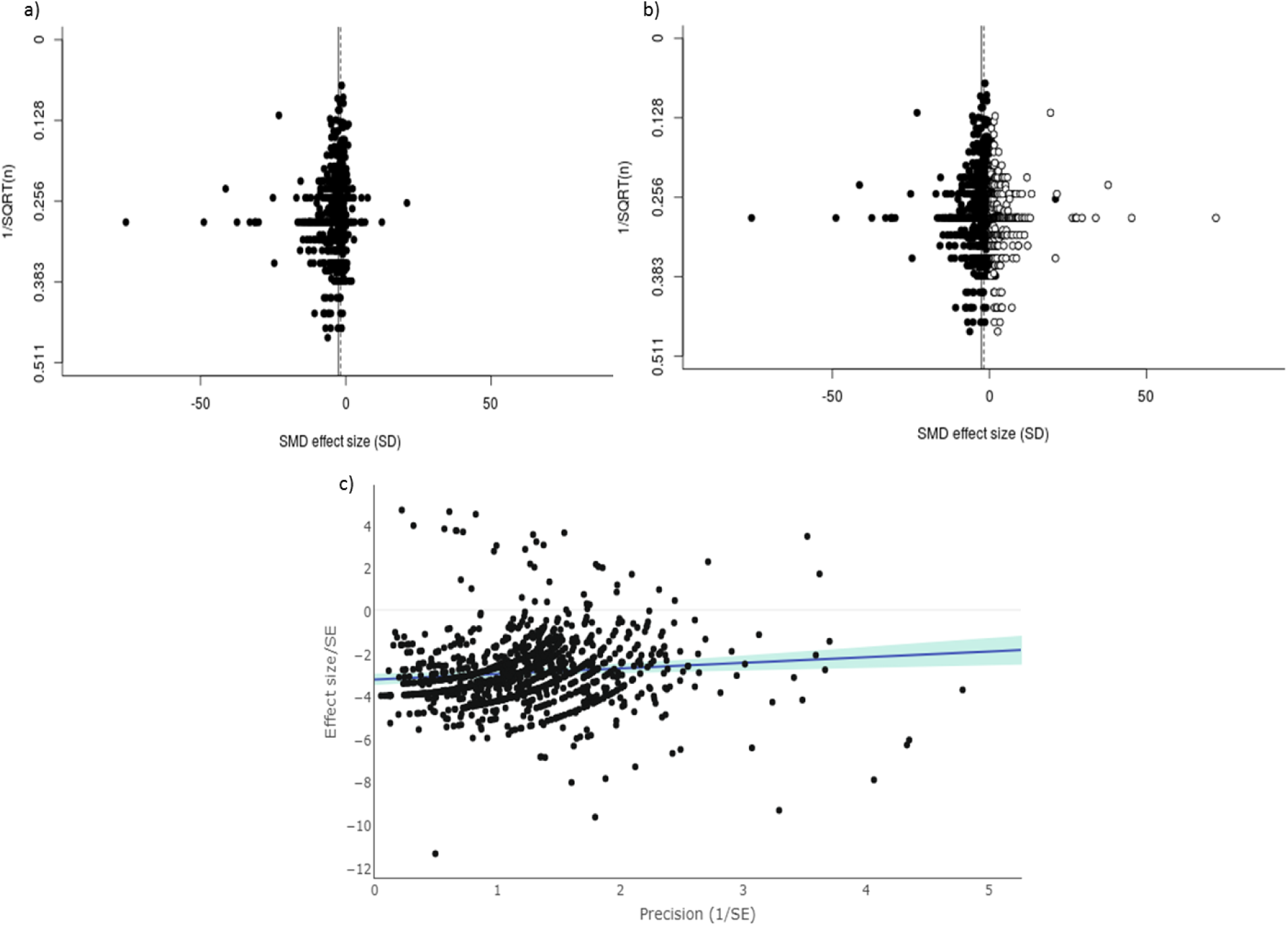
In modelling experiments where a pain-related outcome was used, a) visual inspection of the funnel plot suggests asymmetry. Filled circles represent reported experiments. Solid line represents global effect size and dashed line represents adjusted global effect size. b) trim and fill analysis imputed theoretical missing studies (unfilled circles). Filled circles represent reported experiments. Solid line represents global effect size and dashed line represents adjusted global effect size. c) Egger’s regression indicated small study effects.

For modelling experiments where other behavioural outcome measures were used, there were 88 individual comparisons (−0.71 SD [-0.96;-0.47 CI]). Visual inspection of funnel plots indicated asymmetry, suggesting theoretical missing studies (Figure 12a). Trim and fill analysis imputed 21 theoretical missing studies on the right hand side of the funnel plot, resulting in a total of 109 individual comparisons, Figure 12b. Inclusion of these theoretical missing studies reduced the extent of modelling-induced pain-related behaviour by 56% to - 0.31 SD [-0.58;-0.05]. Further, Egger’s regression indicated small study effects, suggesting funnel plot asymmetry, Figure 12c.

**Figure 12.**
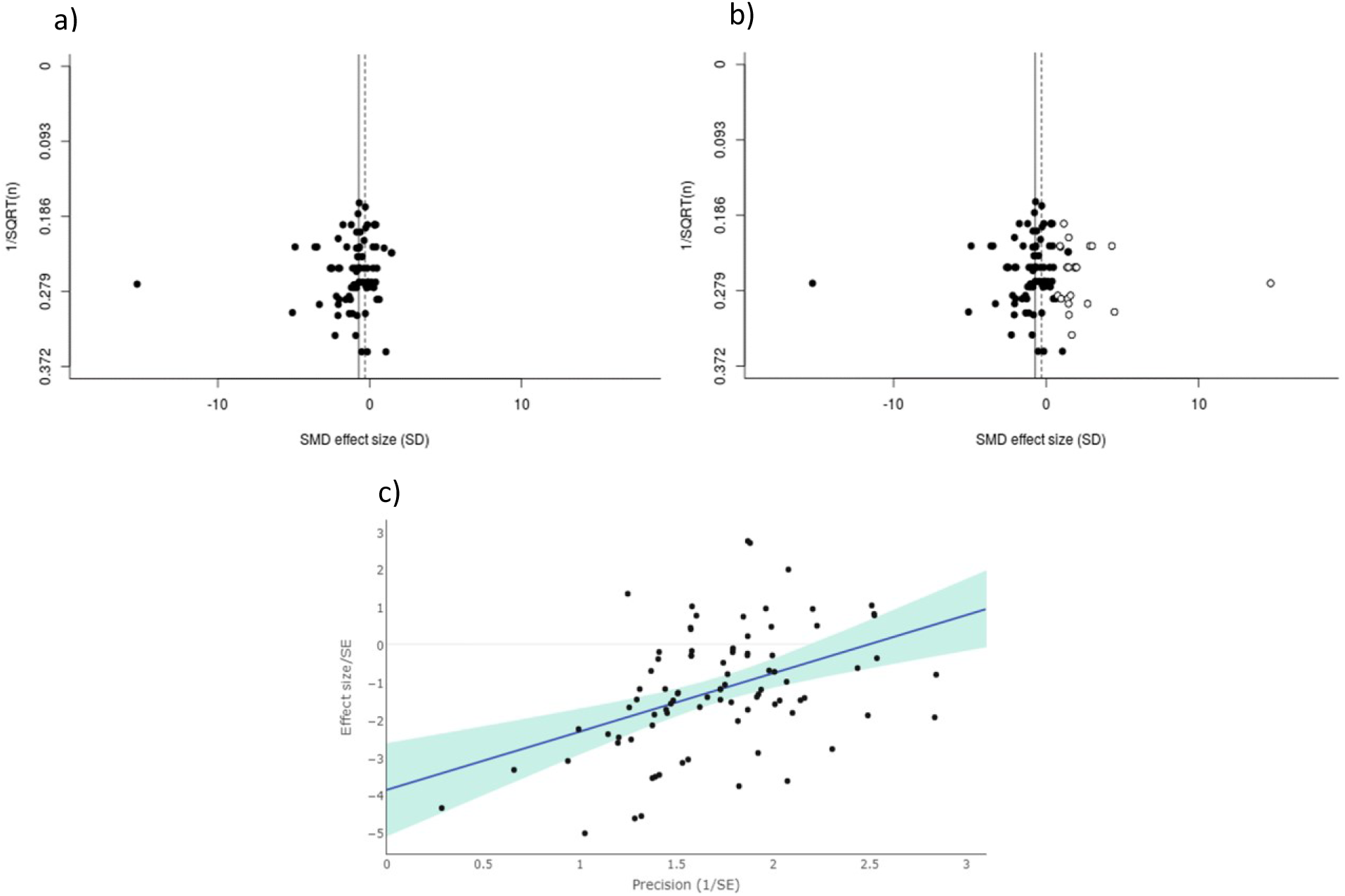
In modelling experiments where other outcomes were used, a) visual inspection of the funnel plot suggests asymmetry. Filled circles represent reported experiments. Solid line represents global effect size and dashed line represents adjusted global effect size. b) trim and fill analysis imputed theoretical missing studies (unfilled circles). Filled circles represent reported experiments. Solid line represents global effect size and dashed line represents adjusted global effect size. c) Egger’s regression indicated small study effects.

## Intervention Experiments

### Drug interventions in Animal Models of CIPN: Pain-Related Behavioural Outcome Measures

In CIPN intervention studies using pain-related behavioural outcome measures, administration of interventions led to a 1.53 SD (1.45; 1.60 95% CI) attenuation of pain-related behaviour compared to control (n=1376 comparisons, p<0.0001). Species did not account for a significant proportion of the heterogeneity (p=0.07), and therefore mice and rat experiments were analysed together.

In intervention studies, the most commonly reported pain-related outcome measure was evoked limb withdrawal to mechanical stimuli (765 individual comparisons), most frequently assessed using mechanical monofilaments to induce limb withdrawal (604 individual comparisons).

### Impact of study design

Intervention accounted for a significant proportion of the heterogeneity, (p=0).The most commonly tested interventions were morphine (n=56 comparisons), gabapentin (n=51 comparisons) and pregabalin (n=39 comparisons) (Figure 13). No clear dose-response relationship was observed for these drug interventions.

**Figure 13.**
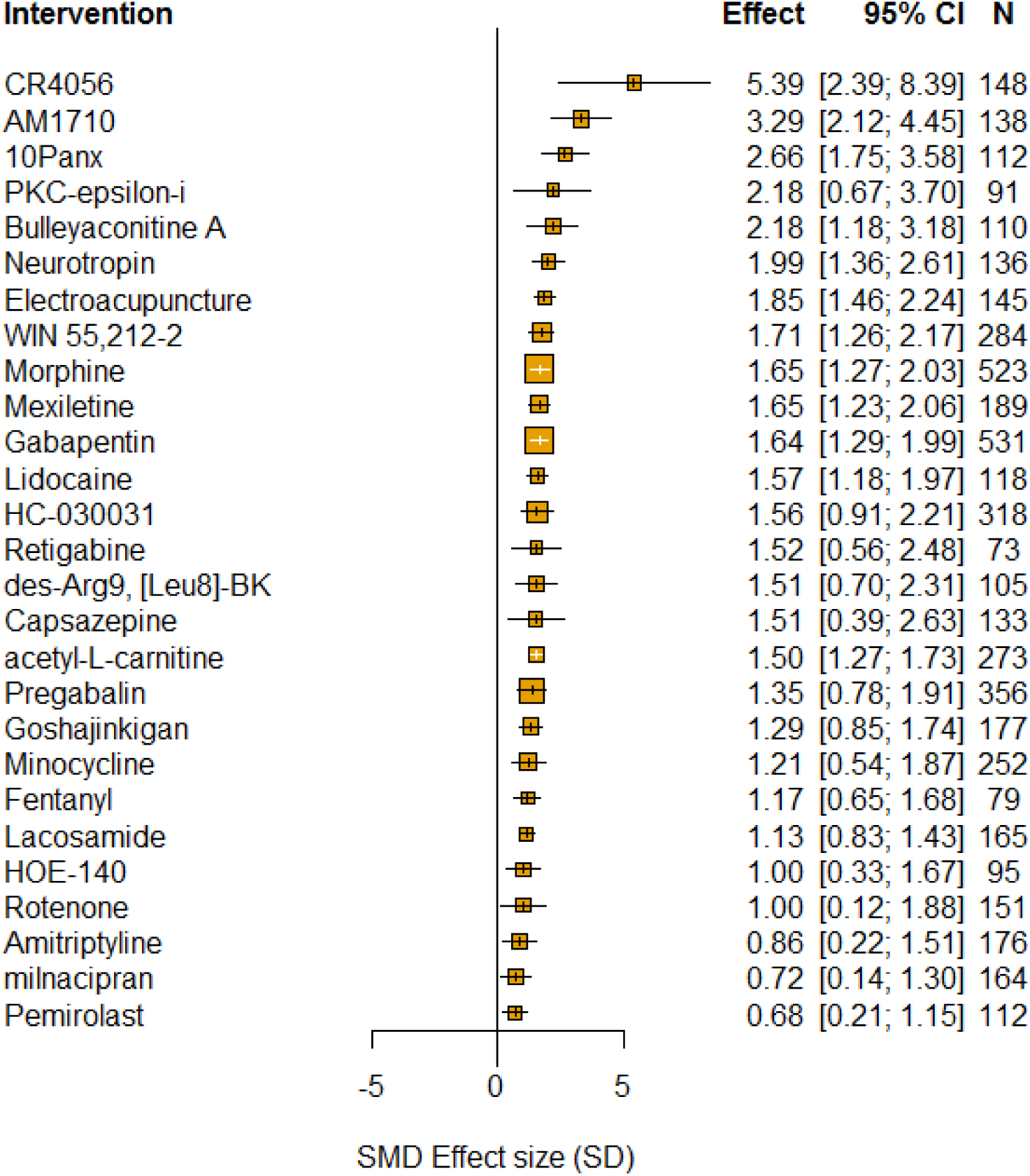
Intervention accounted for a significant proportion of the heterogeneity in intervention experiments using pain-related behavioural outcomes. Plot shows interventions with greater than 10 comparisons. The size of the squares represents the number of nested comparisons that contribute to that data point and the value N represents the number of animals that contribute to that data point.

Pain-related outcome measures accounted for a significant proportion of the heterogeneity. The greatest attenuation of pain-related behaviour was reported by studies that used evoked limb withdrawal to dynamic mechanical touch stimulus (2.36 [-9.71;14.43], n=19, p=6.0×10^−04^, Figure 14a).

**Figure 14.**
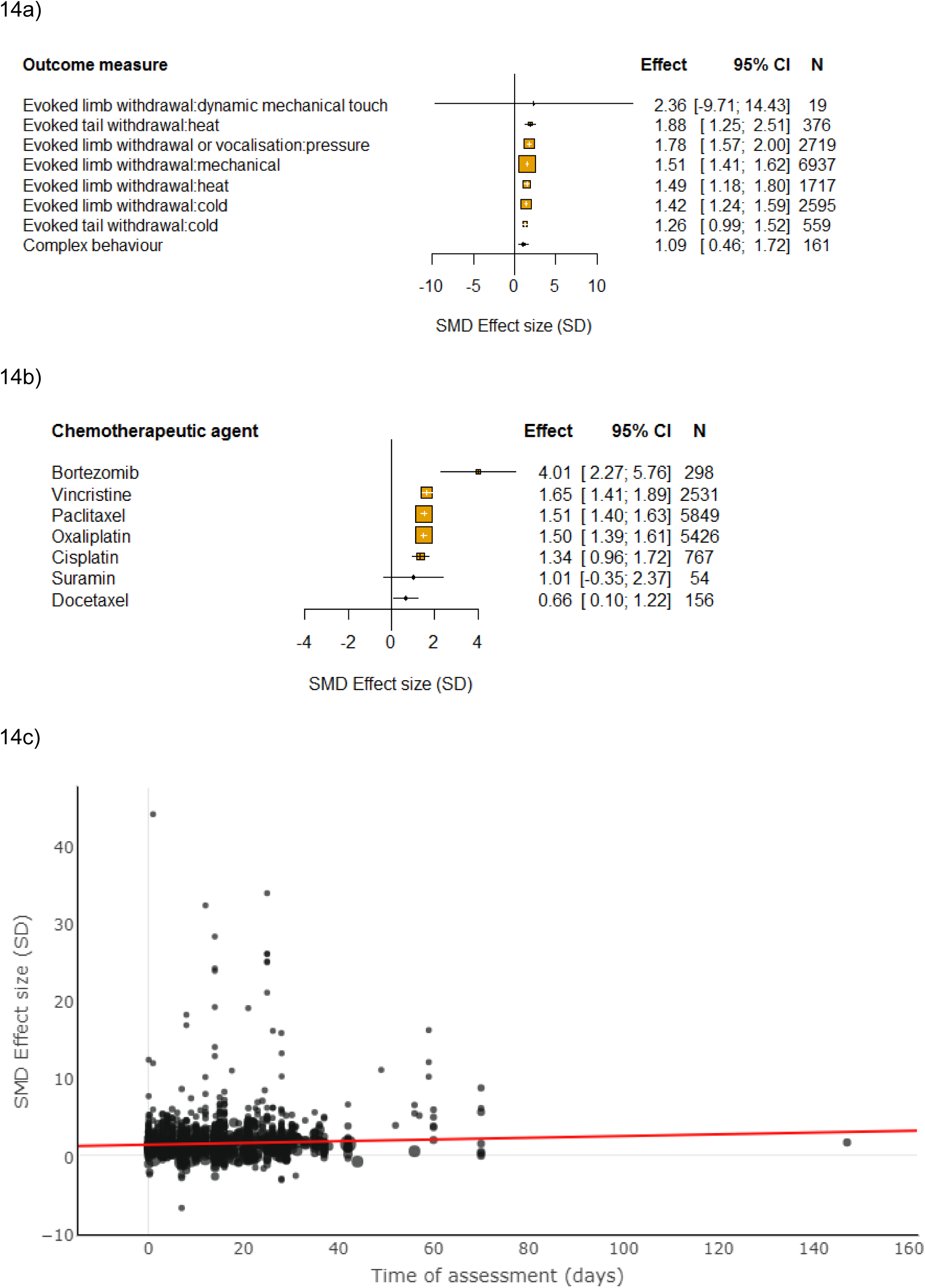
Impact of study design in intervention experiments using pain-related behavioural outcomes. The size of the squares represents the number of nested comparisons that contribute to that data point and the value N represents the number of animals that contribute to that data point. a) Outcome measure accounted for a significant proportion of the heterogeneity. b) Chemotherapeutic agent accounted for a significant proportion of the heterogeneity. c) Time of assessment accounted for a significant proportion of the heterogeneity.

In intervention studies, the chemotherapeutic agent used accounted for a significant proportion of the heterogeneity, with bortezomib associated with the greatest attenuation of pain-related behaviour (4.01 SD [2.27;5.76], n=21, p=1.1×10^−03^, Figure 14b). The most commonly reported were paclitaxel (n=532) and oxaliplatin (n=484).

Sex of animal did not account for a significant proportion of the heterogeneity (p=0.35).

Time to assessment accounted for a significant proportion of the heterogeneity, with a longer interval associated with greater attenuation of pain-related behaviour (p=1×10^−03^, Figure 14c). However, time of intervention administration did not account for a significant proportion of the heterogeneity, p=1.

A *post hoc* analysis found strain of animal accounted for a significant proportion of the heterogeneity (p=0, Figure 15). The most commonly reported were Sprague Dawley rats (n=763).

**Figure 15.**
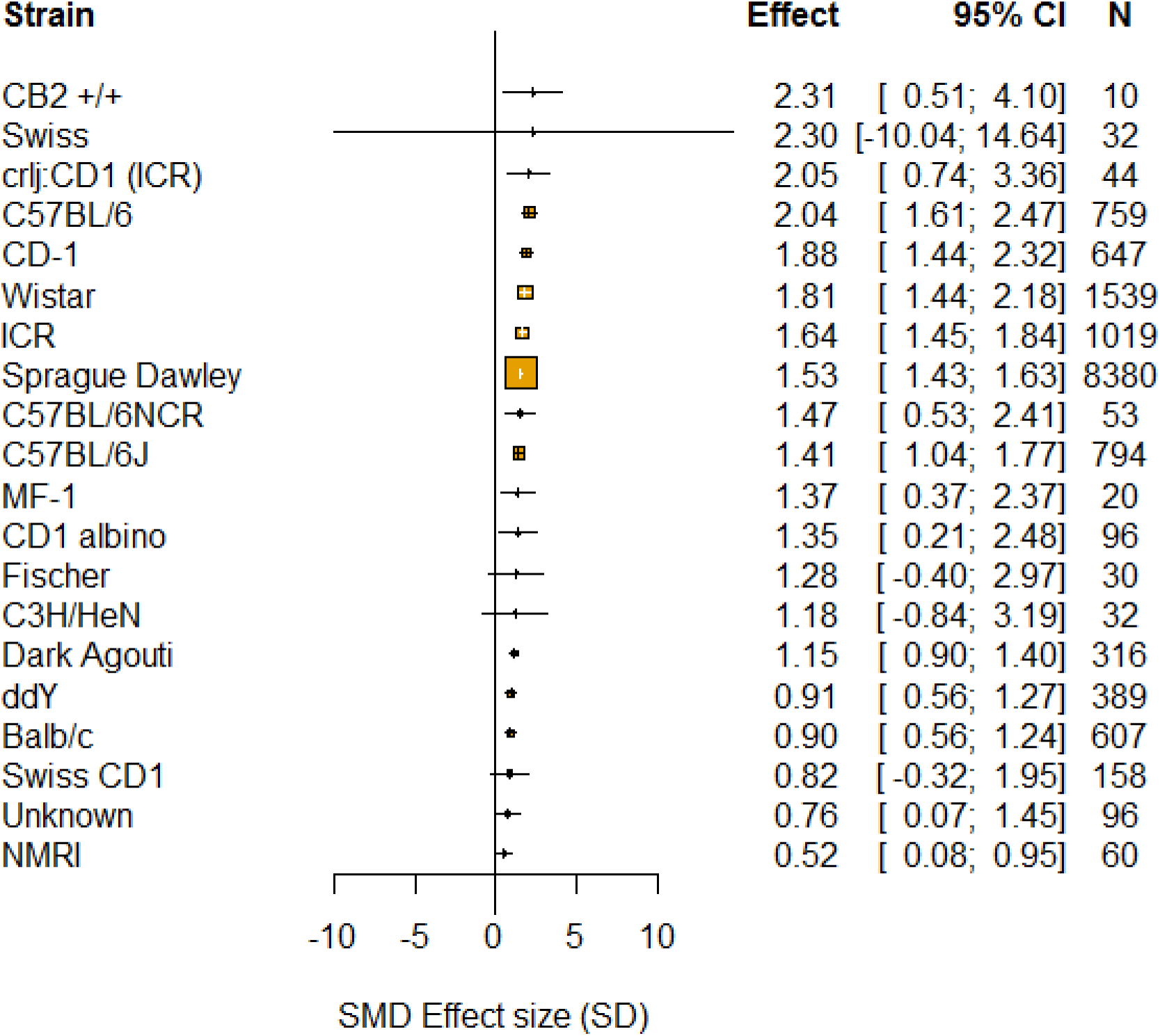
In intervention experiments using pain-related behavioural outcomes, strain accounted for a significant proportion of the heterogeneity. The size of the squares represents the number of nested comparisons that contribute to that data point and the value N represents the number of animals that contribute to that data point.

### Ranking of drug efficacy

A *post hoc* analysis was conducted to compare the ranking of drugs common between a clinical systematic review [15] and our preclinical systematic review. In the clinical systematic review, NNTs were reported for 28 drugs, covering drug class or individual drug. Seventeen of these drugs were also reported by animal studies included in our preclinical systematic review. A Spearman’s rank correlation coefficient found no correlation between clinical and preclinical rank, (r_s_ = -0.0446, p = 0.8652, Table 7 and Figure 16).

**Table 7.**
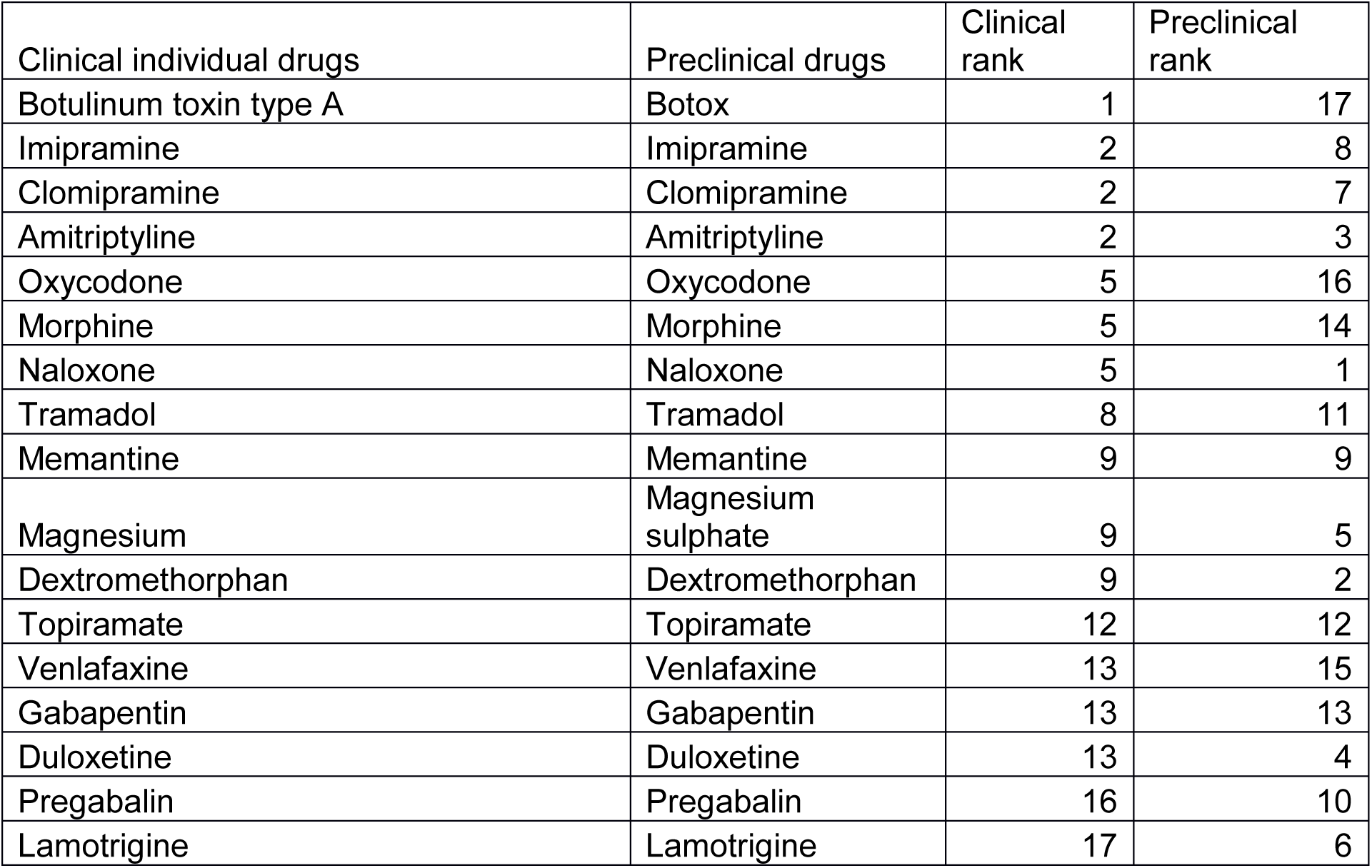
Rank order of drugs in clinical and preclinical meta-analyses.

**Figure 16.**
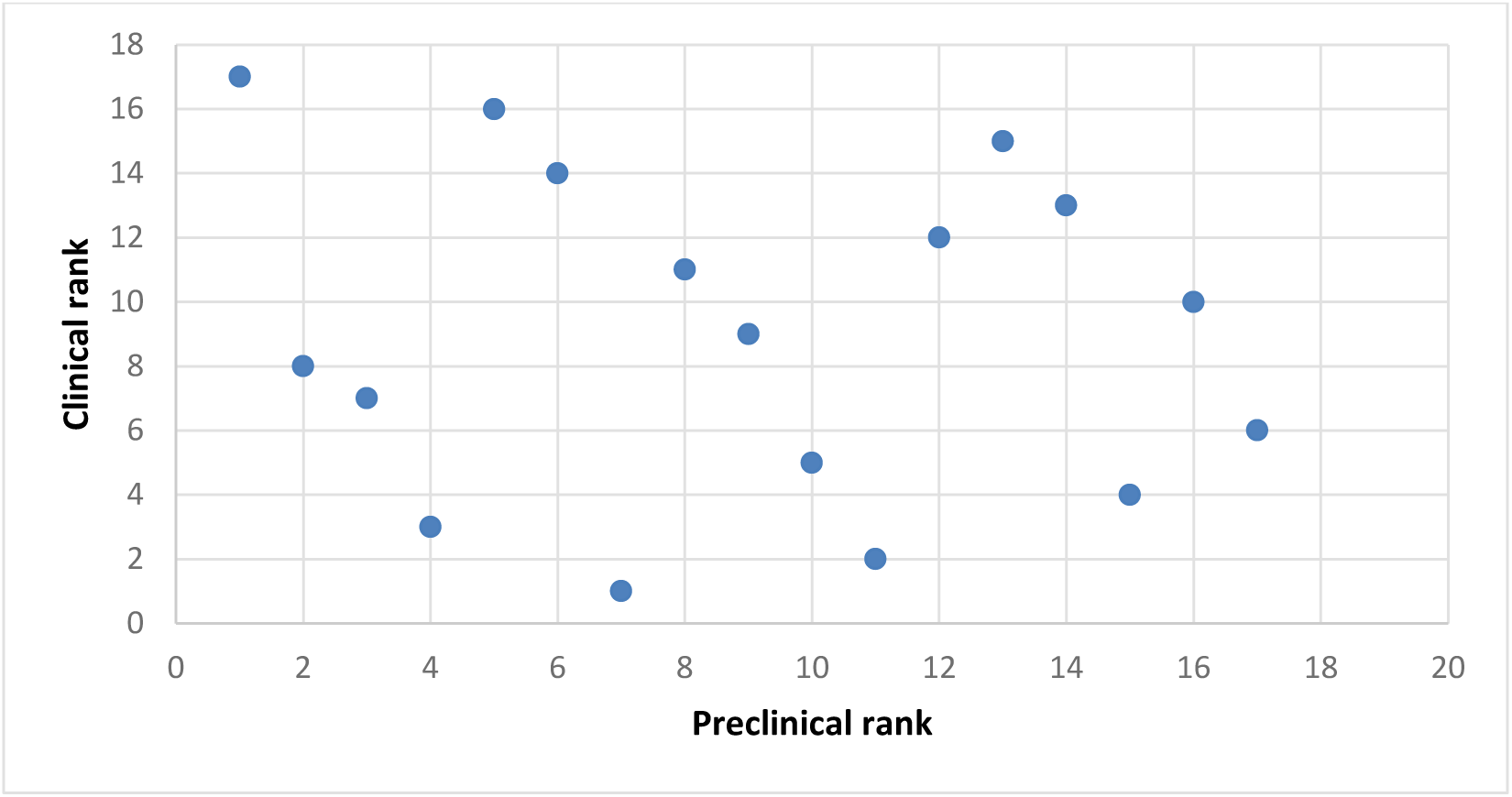
Rank order of clinical and preclinical drugs. A Spearman’s correlation was run to assess the relationship between clinical and preclinical rank of 17 drugs. There was no correlation between clinical and preclinical rank, r_s_ = -0.0446, p = 0.8652.

### Drug Interventions in Animal Models of CIPN: Other Behavioural Outcomes

In intervention studies using other behavioural outcomes, administration of interventions lead to attenuation of other behaviour compared to controls (0.68 SD [0.37;0.99 95% CI], n=37 comparisons). Species did not account for a significant proportion of the heterogeneity (p=0.39).

### Impact of study design

Intervention also accounted for a significant proportion of the heterogeneity, with fentanyl showing in the greatest attenuation of altered reward and locomotor function behaviours (1.83 SD [-3.68;7.34 95%CI], n=2 comparisons, p=7.0×10^−05^, Figure 17).

**Figure 17.**
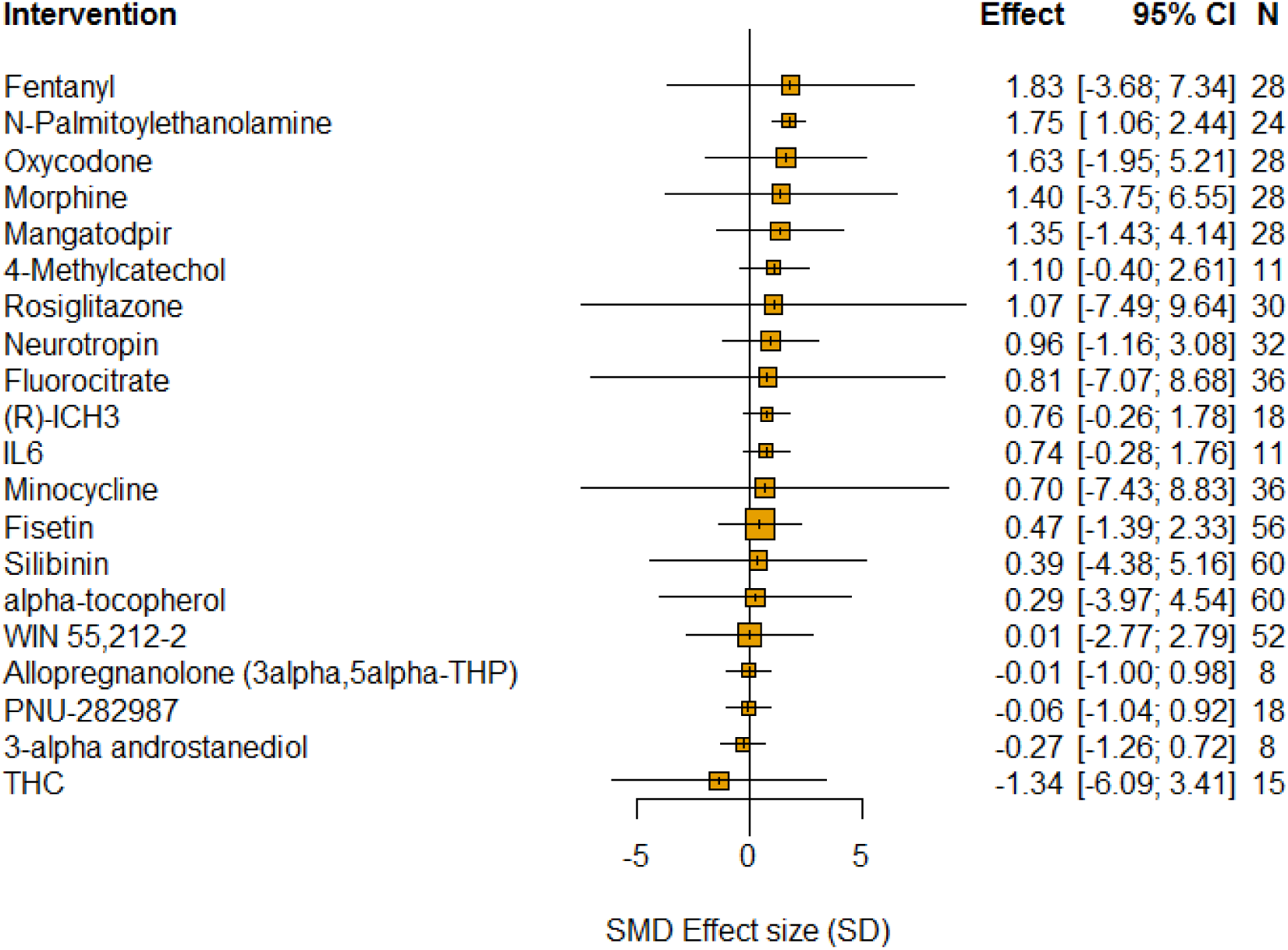
In intervention experiments using pain-related behavioural outcomes, intervention accounted for a significant proportion of the heterogeneity. The size of the squares represents the number of nested comparisons that contribute to that data point and the value N represents the number of animals that contribute to that data point.

Other outcome measures accounted for a significant proportion of the heterogeneity, with reward-related behaviours showing the greatest attenuation by interventions (1.61 SD [1.13;2.07], n=84 comparisons, p=0.0002, Figure 18a).

**Figure 18.**
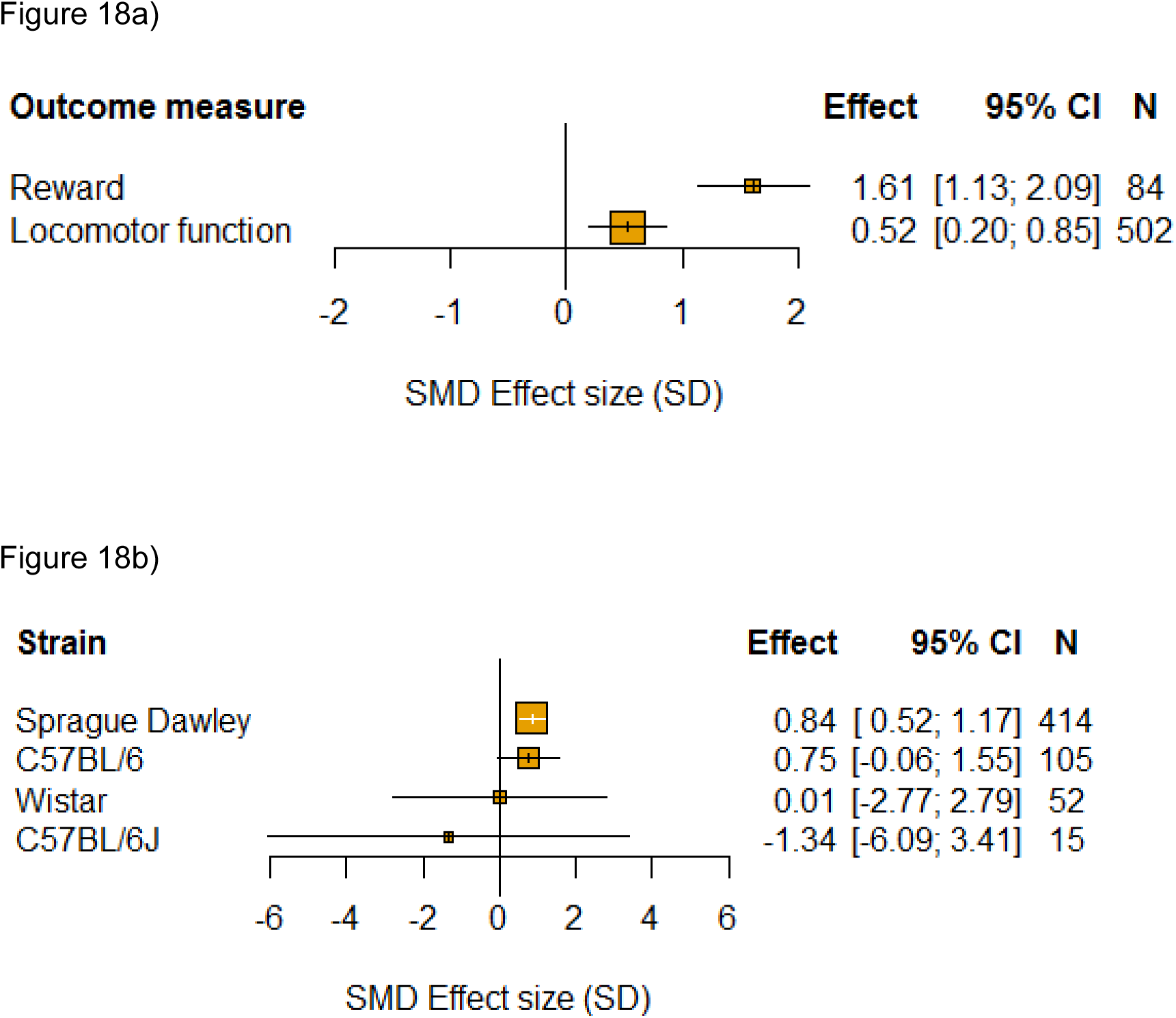
Impact of study design in intervention experiments using other behavioural outcomes. The size of the squares represents the number of nested comparisons that contribute to that data point and the value N represents the number of animals that contribute to that data point. a) Outcome measure accounted for a significant proportion of the heterogeneity. b) Strain accounted for a significant proportion of the heterogeneity.

Chemotherapeutic agent (p=0.137), sex (p=0.008), time to assessment (p=1) and time of intervention administration (p=1) did not account for a significant proportion of the heterogeneity.

In intervention experiments using other behavioural outcome measures, strain account for a significant proportion of the heterogeneity (p=0.001, Figure 18b).

### Power analysis

In intervention studies, the number of animals required to obtain 80% power with a significance level of 0.05 varied substantially across pain-related behavioural tests. For each of mechanical monofilaments, Randall-Selitto paw pressure test, Electronic "von Frey", acetone test/ethylchloride spray, cold plate and Plantar Test (Hargreave’s method), we calculated the number of animals required in model and sham groups. When both mean difference effect size and pooled standard deviation were the 50% centile, the number of animals required ranged from 4 (Randall-Selitto paw pressure test/Hargreave’s thermal test) to 10 (mechanical monofilament), Figure 19. Keeping the mean difference effect size in the 50% centile and increasing pooled standard deviation to the 80% centile increased the number of animals required per group across all behavioural tests and for some tests more than others; from 9 (Hargreave’s) to 104 (mechanical monofilaments). Reducing the mean difference effect size to the 20% centile and pooled standard deviation to the 50% centile, dramatically increased the number of animals required per group, from 21 (Hargreave’s) to 584 (mechanical monofilaments), Figure 19. The values for the 20th, 50th and 80th centile of Mean Differences and pooled standard deviations for each behavioural test are provided (Supplementary Appendix 1).

**Figure 19.**
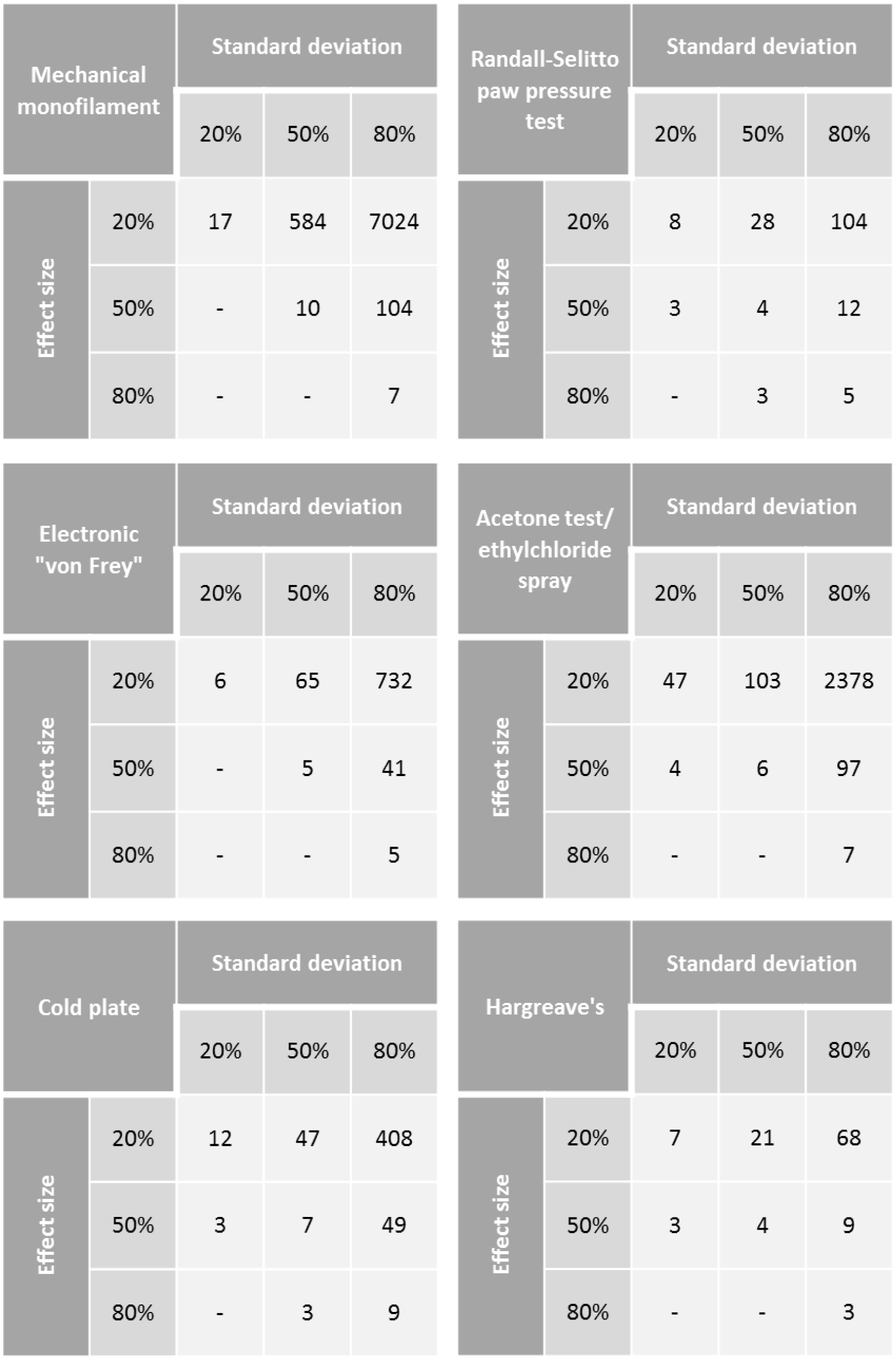
Power analysis for intervention experiments. Number of animals required to obtain 80% power with a significance level of 0.05 using mechanical monofilaments, Randall-Selitto paw pressure test, Electronic “von Frey”, Acetone test/ethylchloride spray, cold plate and Hargreave’s. Effect sizes calculated by Mean Difference. Blank squares could not be calculated.

### Drug interventions in Animal Models of CIPN: Impact of Reporting of Measures to Reduce the Risk of Bias and Measures of Reporting

In CIPN intervention studies using a pain-related behavioural outcome measure, reporting of allocation concealment, animal exclusions and sample size calculations accounted for a significant proportion of the heterogeneity (Figure 9). Failure to report allocation concealment, animal exclusions and sample size calculations were associated with greater estimates of effect. Reporting of randomisation and blinded assessment of outcome did not account for a significant proportion of the heterogeneity.

Both reporting of compliance with animal welfare regulations (p=2.8×10^−03^) and reporting of a conflict of interest statement (p=6.x10^−03^) accounted for a significant proportion of the heterogeneity, (Figure 10). Failure to report this information was associated with decreased estimates of effect (Table 8).

**Table 8.** Intervention comparisons using pain-related behavioural outcome measures. Stratified meta-analysis results for reporting of measures to reduce the risk of bias and measures of reporting (p<0.007).

|  | Intervention comparisons pain-related (n=1376) |  |  |  |  |  |
| --- | --- | --- | --- | --- | --- | --- |
|  | Accounts for a significant proportion of heterogeneity | Not reporting associated with | Reported | Effect | SE | No. of comparisons |
| Risk of bias |  |  |  |  |  |  |
| Blinded assessment of outcome | No | NA | TRUE | 1.491 | 0.05 | 762 |
|  |  |  | FALS<br>E | 1.581 | 0.064 | 614 |
| Random allocation to group | No | NA | TRUE | 1.615 | 0.09 | 383 |
|  |  |  | FALS<br>E | 1.502 | 0.043 | 993 |
| Allocation concealment | Yes | Increased estimate of effect | TRUE | 0.941 | 0.248 | 25 |
|  |  |  | FALS<br>E | 1.541 | 0.04 | 1351 |
| Sample size calculation | Yes | Increased estimate of effect | TRUE | 1.037 | 0.392 | 14 |
|  |  |  | FALS<br>E | 1.534 | 0.04 | 1362 |
| Animal exclusions | Yes | Increased estimate of effect | TRUE | 1.408 | 0.105 | 215 |
|  |  |  | FALS<br>E | 1.552 | 0.042 | 1161 |
| Reporting |  |  |  |  |  |  |
| Compliance with animal welfare regulations | Yes | Decreased estimate of effect | TRUE | 1.534 | 0.04 | 1365 |
|  |  |  | FALS<br>E | 0.514 | 0.217 | 11 |
| Conflict of interest statement | Yes | Decreased estimate of effect | TRUE | 1.638 | 0.061 | 744 |
|  |  |  | FALS<br>E | 1.423 | 0.05 | 632 |

In intervention studies using other behavioural outcome measures, reporting of randomisation, allocation concealment, animal exclusions or sample size calculation did not account for a significant proportion of the heterogeneity. Blinded assessment of outcome accounted for a significant proportion of the heterogeneity (p=0.0044), with those that did not report blinded assessment of outcome were associated with larger effect sizes (Figure 9). Reporting of conflict of interest statement or compliance with animal welfare regulations did not account for a significant proportion of the heterogeneity (Table 9).

**Table 9.** Intervention comparisons using other behavioural tests. Stratified meta-analysis results for reporting of measures to reduce the risk of bias and measures of reporting (p<0.007).

|  | Intervention comparisons other (n=37) |  |  |  |  |  |
| --- | --- | --- | --- | --- | --- | --- |
|  | Accounts for a significant proportion of heterogeneity | Not reporting associated with | Reported | Effect | SE | No. of comparisons |
| Risk of bias |  |  |  |  |  |  |
| Blinded assessment of outcome | Yes | Increased estimate of effect | TRUE | 0.454 | 0.18 | 22 |
|  |  |  | FALSE | 1.113 | 0.242 | 15 |
| Random allocation to group | No | NA | TRUE | 0.417 | 0.305 | 15 |
|  |  |  | FALSE | 0.829 | 0.161 | 22 |
| Allocation concealment | NA | NA |  |  |  |  |
|  |  |  | FALSE | 0.685 | 0.153 | 37 |
| Sample size calculation | NA | NA |  |  |  |  |
|  |  |  | FALSE | 0.685 | 0.153 | 37 |
| Animal exclusions | No | NA | TRUE | 0.561 | 0.222 | 12 |
|  |  |  | FALSE | 0.755 | 0.207 | 25 |
| Reporting |  |  |  |  |  |  |
| Compliance with animal welfare regulations | No | NA | TRUE | 0.685 | 0.153 | 37 |
| Conflict of interest statement | No | NA | TRUE | 0.599 | 0.175 | 21 |
|  |  |  | FALSE | 0.855 | 0.279 | 16 |

### Assessment of publication bias in animal CIPN models that test efficacy of drug interventions

For intervention that used a pain-related outcome measure, there were 1529 individual comparisons (1.52 SD [1.44;1.60 95%CI]). Visual inspection of funnel plots indicated asymmetry, suggesting theoretical missing studies, Figure 20a. Trim and fill analysis imputed 393 theoretical missing studies on the left hand side of the funnel plot, resulting in a total of 1922 individual comparisons, Figure 20b. Inclusion of these theoretical missing studies decreased the estimate of modelling induced pain-related behaviours by 28% to 1.09 SD ([1.01;1.16]). Further, Egger’s regression indicated small study effects, suggesting funnel plot asymmetry, Figure 20c.

**Figure 20.**
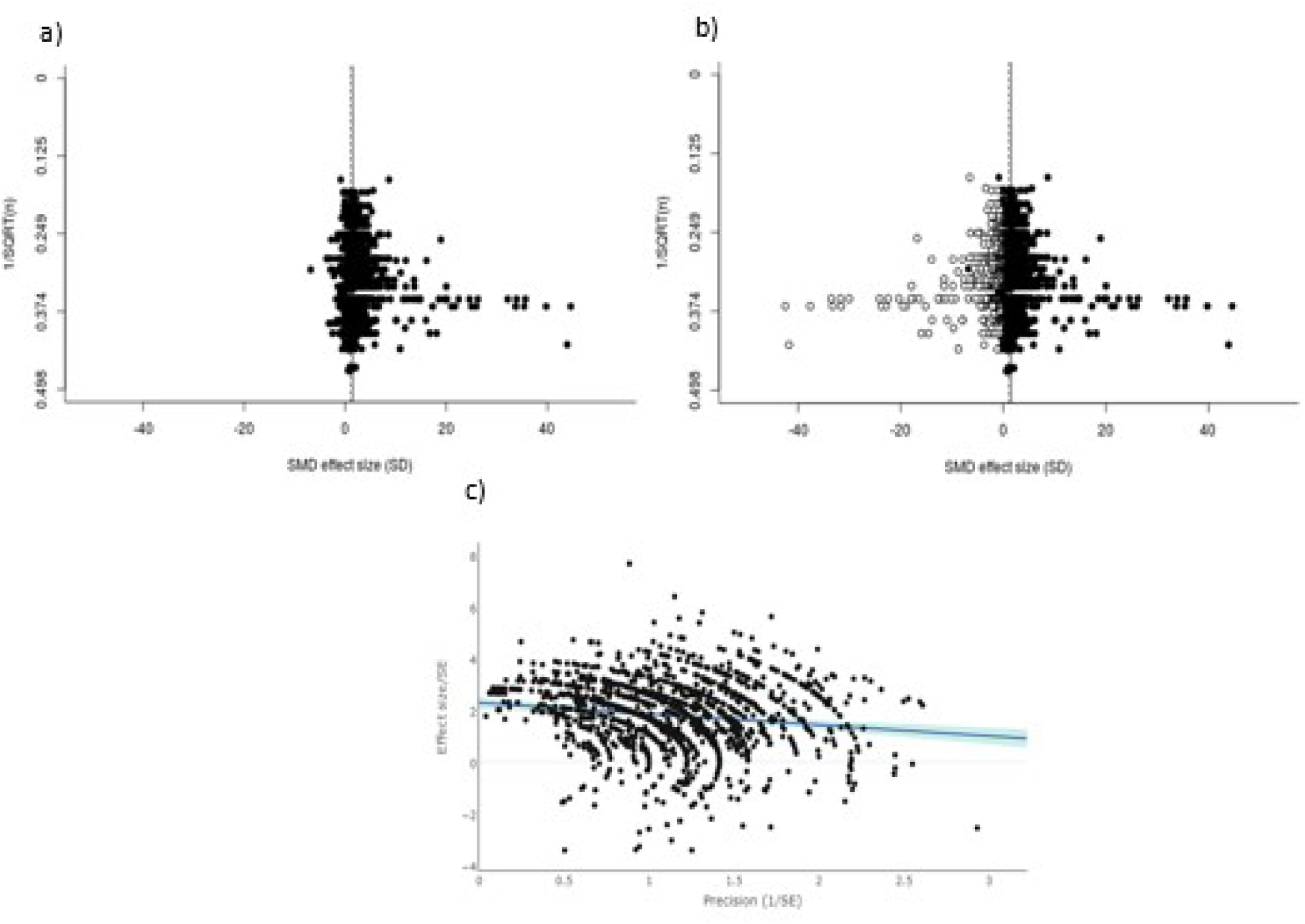
In intervention experiments where a pain-related outcome was used, a) visual inspection of the funnel plot suggests asymmetry. Filled circles represent reported experiments. Solid line represents global effect size and dashed line represents adjusted global effect size. b) trim and fill analysis imputed theoretical missing studies (unfilled circles). Filled circles represent reported experiments. Solid line represents global effect size and dashed line represents adjusted global effect size. c) Egger’s regression indicated small study effects.

For intervention studies that used other behavioural outcome measures, there were 43 individual comparisons (0.70 SD [0.41;0.99 CI]). Visual inspection of funnel plots did not indicate asymmetry suggesting no theoretical missing studies, Figure 21a. Trim and fill analysis estimated no theoretical missing studies on the left hand side of the funnel plot. Further, Egger’s regression indicated small study effects, suggesting funnel plot asymmetry, Figure 21b.

**Figure 21.**
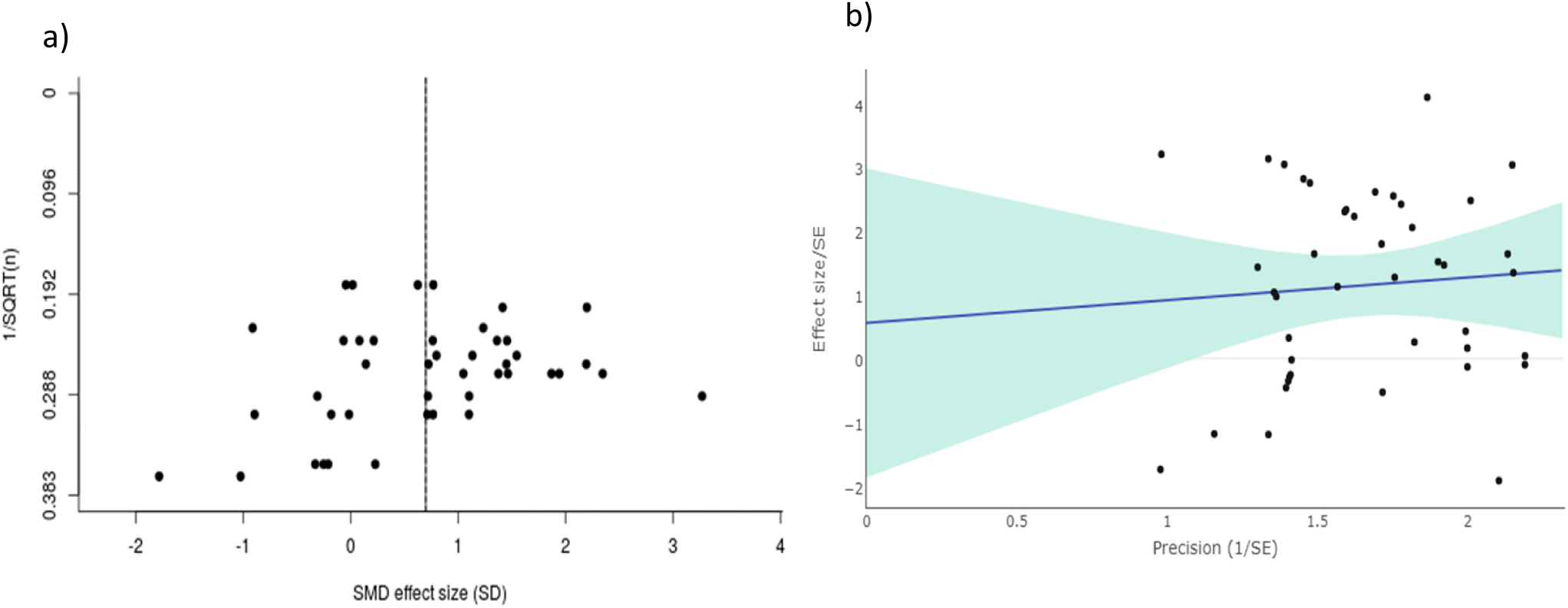
In intervention experiments where other outcome measures were used, a) visual inspection of the funnel plot does not suggest asymmetry. Filled circles represent reported experiments. Solid line represents global effect size and dashed line represents adjusted global effect size. b) Egger’s regression indicated small study effects.

### Animal husbandry

The reporting of animal husbandry details was low across all included studies (Supplementary Table 1).

## Discussion

The results of our systematic review indicate that the publication rate for reports of experiments using animal models of CIPN is high. Between the initial search in 2012 and the updated search in 2015, the number of relevant publications doubled: 181 relevant publications were identified in 2012, and a further 161 relevant publications were identified 3 years later in 2015. The high publication accrual rate is not unique to this field but is the case across clinical [7] and preclinical research; making it challenging for researchers and consumers of research to keep up-to-date with the literature in their field. This is the first systematic review of preclinical models of pain to use machine-learning and text mining, and the first to demonstrate the usefulness of these automation tools in this field.

### External validity of studies using animal models of CIPN Misalignment between animal models and the clinical population

In the clinic, the chemotherapeutic agents included in this analysis are frequently used to treat women cancer patients. However the majority of studies (285/341 (84%)) identified by this systematic review used only male animals to model CIPN, reducing the external validity of the findings from these models to the clinical population.

We must consider whether we are modelling the same condition as that observed in the clinic. Acute CIPN is estimated to affect 68.1% of patients within the first month of chemotherapy cessation, 60% at 3 months, and 30% of patients at 6 months [39]. Severe acute CIPN may require dose reduction or cessation of chemotherapy [5] however many patients symptoms improve after chemotherapy cessation. Taxane chemotherapeutic agents led to CIPN in up to 80% of exposed patients 2 years after treatment [18] and oxaliplatin chemotherapy led to peripheral neurotoxicity in 79.2% of patients at 25 months post treatment [34]). A long-term study into oxaliplatin showed treatment was associated with peripheral neurotoxicity at 6 years follow-up [25]. Therefore, the models of CIPN identified in this systematic review likely model the acute phase of CIPN. Of those publications where model duration (the time between the first administration of chemotherapeutic agent and the time when all animals were killed) was reported (39 of 341 publications), the median model duration was 21 days [16-28 IQR]. Further, the median time to assessment in our modelling dataset (the time where there was the biggest difference between CIPN model and sham animals), was 14 days [7-25 IQR]. Interestingly, the median time to assessment in our intervention dataset was 14 days [7-22 IQR]. Indicating that the time where the drug interventions are most effective is when the models show the largest modelling effect.

### Misalignment between preclinical and clinical outcome measures

The most frequent behaviours reported in animal models of chemotherapy-induced peripheral neuropathy are estimates of gain in sensory function; hypersensitivity in paw withdrawal evoked by mechanical stimuli was the assay most often employed. A review of studies reporting preclinical models of pain published in the Journal of Pain between 2000 and 2004, showed that the most commonly reported pain-related behaviours were also reflex withdrawal responses [31]. This starkly contrasts with chronic clinical CIPN, where the predominant clinical sensory phenotype of these patients is one of sensory loss [44]-i.e. the opposite of that reported in animal models. There is thus a dichotomous mismatch between the sensory profile reported from animal studies and what is observed in chronic CIPN patients, potentially compromising the clinical relevance of these models for chronic CIPN, however, they may have more relevance to acute CIPN.

To address the misalignment between outcome measures used to assess pain in patients in clinical trials and those commonly reported in animal models of CIPN, we have called for the development of sensory profiling for rodent models of neuropathy that reflect the clinical methods [35].

Others have shown that external validity can be increased by using multi-centre studies to create more heterogeneous study samples, an approach that may be useful in pain modelling [46]

### Internal validity of studies using animal models of CIPN

There was moderate reporting of measures to reduce the risk of bias. Statistical modelling and meta-analysis have demonstrated that the exclusion of animals can distort true effects, where random loss of samples decreased statistical power and biased removal dramatically increased the probability of false positive results [20]. It has been shown in other research fields that efficacy is lower in studies that report measures to reduce the risk of bias [8; 28; 36; 45].

The details of methods used to implement randomisation and blinding and the methods and assumptions for sample size calculation were rarely reported. These are important to understand the quality of these procedures, as opposed to mere reporting. If methods and assumptions were reported this would allow assessment of the quality of these procedures using a tool such as Jadad scoring [24].

### Publication bias

Publication bias analysis suggested that taking the many imputed missing studies into account dramatically reduced the global effect sizes in all data sets, except the smallest data set (n=37). Publication bias is a known problem in preclinical research, where studies with small effect sizes or where no (or one opposing that hypothesised) effect is observed are likely to remain unpublished whilst positive findings prioritised. Others have demonstrated the presence of publication bias in preclinical research [38], One potential reason for this is the high competition for promotion, funding and publishing that results in few incentives to spend time publishing results from studies where null hypothesis was not disproved. We would like to see more incentives, such as Registered Reports for publishing well designed, thoroughly reported studies, regardless of the results.

### Optimising experimental design

Experimental design of *in vivo* CIPN studies can be optimised by adopting measures to reduce the risk of bias, such as using power calculations to ensure experiments are appropriately powered. It is also imperative to use a model that best represents the clinical population of interest, for example using both female and male animals. To help further address the issue of publication bias, we suggest that researchers make protocols for preclinical studies available and publish all results.

One approach that would help optimise experimental design is to use the Experimental Design Assistant (https://eda.nc3rs.org.uk/), a free resource developed by the NC3Rs, whereby researchers create a record of their experimental design. This could then be uploaded to the Open Science Framework as a record.

### Reduction

It may be that we see a shift towards reducing waste and maximising the information gained from our studies. This would require open and transparent reporting of results. For complex behaviours, even if the tests were not primarily performed to test for pain-related behaviour, the online publication as supplementary material of individual animal video files [33], would allow re-analysis for pain-related behaviours such as thigmotaxis. Future studies are encouraged to publish such files to allow secondary analyses of complex behaviours and thereby reduce the number of animals used in future studies. It is interesting to note that no publication in this systematic review of animal models of CIPN used open field to measure thigmotaxis, as is reported in other preclinical pain research.

Our exemplar power calculations of the most commonly reported behavioural outcome measures highlight the substantial variability in the statistical performance of different outcome measures. Using these results it is possible to rank the different pain-related behavioural tests according to how many animals are required per group if effect size or standard deviation increases or decreases. Along with other factors, such as clinical relevance, these results can inform the choice of outcome measure in study design, by allowing researchers to select outcomes measures that minimise the number of animals required.

### Limitations

Conducting a systematic review is time and resource-intensive, and the rate of publication means systematic reviews rapidly become outdated. This review is limited as the most recent information included was identified in November 2015. We propose that the present systematic review form a “baseline” systematic review, which can be updated and developed into a living systematic review, i.e. a systematic review that is continually updated as new evidence becomes available [15]. As new online platforms and tools for machine-learning and automation become available, preclinical living systematic reviews become more feasible [42]. Living systematic review guideline have recently been published [14] and Cochrane have also launched pilot living systematic reviews [3; 41].

The machine-learning algorithm based on our initial screening had a high sensitivity (97%) and medium specificity (67%). High sensitivity captures most relevant literature and has a low risk of missing relevant literature. An algorithm with lower specificity is more likely to falsely identify included studies (i.e. more irrelevant studies are identified as included), compared to an algorithm with high specificity. With a specificity of 67%, our algorithm did falsely identify some irrelevant studies, which resulted in the two independent human screeners having more studies to exclude when it came to data abstraction. We believe that this balance between sensitivity and specificity was appropriate as this reduced the risk of missing relevant studies.

In our meta-analysis, we grouped together the behavioural outcome measures that measure the same underlying biology. For example, in the case of experiments that reported using the grip test; five studies reported that the test was used to measure grip strength, and one reported that the test was used to measure muscle hyperalgesia [23]. For this reason, in our analysis we grouped all grip test outcome measures together as a non-pain-related behavioural outcome measures. It is possible that the same tests or similar tests could be used and the same measurements reported as different outcomes. This is one of the challenges when analysing published data.

We only included studies where the intervention drug was administered after or at the same time as the chemotherapeutic agent. Future literature reviews may look at those drug interventions given before chemotherapeutic agents to determine if drug intervention at this time point can effectively prevent CIPN.

Unfortunately, the reporting of measures to reduce the risk of bias was moderate in the studies included in this systematic review, which limits what we can infer from the results. We hope this review will act to highlight this issue in the field of CIPN *in vivo* modelling. Systematic review of animal experiments in other research areas has revealed low reporting of these measures and negative impact of failure to report these measures, across *in vivo* fields as diverse as modelling of stroke, intracerebral haemorrhage, Parkinson’s disease, multiple sclerosis and bone cancer pain [9; 17; 30; 36; 37; 45] This has driven change, influencing the development of reporting guidelines [26], pain modelling specific guidelines [4] and the editorial policy of Nature Publishing Group [1]. Encouragingly, after an initial review on the efficacy of Interleukin-1 Receptor antagonist in animal models of stroke highlighted low reporting of measures to reduce the risk of bias [6], a subsequent review identified increased reporting of these measures [30], increasing the validity and reliability of these results. We anticipate that there will be a similar improvement in studies reporting the use of animal models of CIPN. We propose that once studies achieve sufficient quality, it will be possible to use a GRADE type analysis or process to rate the certainty of the evidence of animal studies [21]. The measures to reduce the risk of bias that we have assessed for the reporting of are largely derived from what is known to be important in clinical trials, and the extent to these measures impact upon the findings of animal studies has yet to be fully elucidated. However, reporting of these measures allows users of research to make informed judgment about findings. Finally, it should be noted that there are likely to be other measures that are important in animal studies that we have not considered.

## Conclusions

This systematic review and meta-analysis provides a comprehensive summary of the *in vivo* modelling of CIPN with the aim of informing robust experimental design in future studies. Areas where the internal and external validity preclinical CIPN studies can be increased have been identified. The external validity can be increased by using both sexes of animals in the modelling of CIPN and ensuring outcome measures align with those most relevant in the clinic. Measures to reduce the risk of bias should be employed to increase the internal validity of studies. Power analysis calculations illustrate the variation in group size under different conditions and between different behavioural tests, and can be used to inform outcome measure choice in study design.

## Acknowledgements

This work is part of the NC3Rs funded project; Reduction and refinement in animal models of neuropathic pain: using systematic review and meta-analysis. This work is also part of the Europain Collaboration, which has received support from the Innovative Medicines Initiative Joint Undertaking, under grant agreement no 115007, resources of which are composed of financial contribution from the European Union’s Seventh Framework Program (FP7/2007-2013) and EFPIA companies’ in kind contribution.

Reporting of animal husbandry details. The median habituation time was 7 days [7-7 IQR]. The median number of animals per cage was 4 [2.5-4.5]. Reporting of mixed housing with shams was always ‘Not mixed’. Room temperature 22’C [22-23 IQR]. Humidity 55 [53.75-55 IQR].

